# A transient pulse of neuronal ATF4 installs a durable non-autonomous programme promoting healthy ageing

**DOI:** 10.64898/2026.11.02.763634

**Authors:** Paula Suárez-Distasio, Óscar Tovar-Ferrero, Sasha Saint-Roman, Oliver Mitev, Marc Liesa, Guillermo Martínez Corrales

## Abstract

Stresses occurring transiently and early in life can have long-standing consequences for health, even long after the stress has resolved. Many environmental stressors converge on activating the integrated stress response (ISR), whose major effector is ATF4. However, it remains unknown whether transient ATF4-ISR activation early in life is a mediator of life and healthspan. Here we used inducible, reversible ATF4 gain-of-function in *Drosophila* to determine how the timing, duration and tissue of ATF4 activation determine ageing outcomes. We found that ATF4 induction confined to early adulthood extended healthspan, reducing mortality risk by approximately 50% across all ages and improving neuromuscular function in aged flies. Multi-organ transcriptomics showed that the acute ATF4 transcriptional response was transient, as expected, but replaced by a distinct, secondary transcriptional programme of metabolic remodelling that was associated with decreased total body protein content. By using tissue-specific ATF4 induction, we found that transient ATF4 activation in neurons was sufficient to extend lifespan, slow age-related locomotor decline and increase starvation resistance in old age. This neuron-restricted activation prevented the costs to body weight and fecundity induced by ubiquitous ATF4 activation. Transient neuronal ATF4 left a lasting non-autonomous imprint on the fat body at old age: induction of heterochromatin genes, reduced transposable element expression, and increased mitochondrial respiratory capacity. These findings show that the duration and tissue of ISR-ATF4 activation are the key determinants of ATF4 actions that promote healthy ageing, while revealing a brain-to-fat body axis initiated by neuronal ATF4 that results in lasting systemic adaptation.

## Introduction

Early-life experiences can have profound and lasting consequences for health and survival in later life. In humans, adverse conditions in childhood, such as poor nutrition or stress, are associated with altered health and survival decades later, effects that can persist even across generations ^1–3^. Experimental work in model organisms has shown that early-life activated programming can be beneficial as well: interventions confined to a brief window of early adulthood can produce health benefits that last long after the intervention has ended. For example, in *Drosophila*, brief rapamycin treatment in early adulthood extends lifespan as effectively as lifelong treatment^4^, and we showed that transient activation of the transcription factor dFOXO in young flies reduces mortality later in life through lasting chromatin remodelling^5^. These findings suggest that the ageing trajectory can be set early in life by the transient activity of signalling and stress-response pathways. Which pathways can be programmed in this way, in which tissues, and for how long they need to be active remain largely unknown.

One candidate pathway is the integrated stress response (ISR), a conserved pathway through which cells sense diverse environmental stresses, including amino acid deprivation, endoplasmic reticulum and mitochondrial stresses^6,7^. The ISR adjusts gene expression and protein translation to ensure survival and resistance to stress. Its main transcriptional effector, ATF4, drives programmes of amino acid metabolism, antioxidant defence and protein folding that support survival. However, sustained and strong ATF4 activation caused by unresolved stresses cause cell death^8^. Consistent with this duality, the role of ATF4 in ageing and disease remains contradictory. ATF4 orthologues extend lifespan when overexpressed in yeast and worms^9,10^, and ATF4 activity is increased in several long-lived mouse models^11^. Conversely, loss of ISR signalling also extends lifespan in worms^12^, and ATF4 promotes muscle ageing in different contexts ^13,14^.

These studies point to the tissue in which ATF4 is activated, as well as inter-organ communication might be the key aspects determining ATF4-ISR mediated benefits or pathology. In *C. elegans*, mitochondrial stress or activation of the unfolded protein response in neurons is sufficient to induce protective responses in distant tissues and extend lifespan^15,16^. Stress-response transcription factors can also act through different programmes in different tissues, as shown for Xbp1 in flies^17^. Even within the same tissue and in response to a similar insult, ATF4 itself can be harmful or protective: in the fly brain, ATF4 has been reported to both cause neurodegeneration^18^ to prevent it^19,20^, yet its effect on organismal ageing when activated specifically in neurons has not been examined. Indeed, neuron-specific transient ATF4 activation has not been tested for its effects on lifespan in any organism, leaving open whether a brief ISR in a single tissue, such as the nervous system, could programme the ageing of the whole animal.

By using multi-organ transcriptomics and high resolution respirometry in an array of tissue-specific and inducible *in vivo* ageing *Drosophila* models, we find that early adulthood, transient ATF4 activation extends lifespan at a cost to body weight and fecundity, while chronic ATF4 activation in adulthood shortens lifespan. Multiple models of activation restricted to individual organs revealed that activation in neurons, but not in the fat body, gut or muscle, is sufficient to extend lifespan and slow locomotor decline, at no cost to reproduction or body weight. This brief neuronal pulse leaves a lasting imprint on a distant tissue: weeks after the stimulus has ended, the fat body of old flies shows induction of heterochromatin genes, a downward shift in transposable element expression and remodelled mitochondrial respiration. Our findings identify neurons as a site where a transient stress response early in adult life is converted into durable, organism-wide adaptations that promote healthy ageing.

## Methods

### Reagents and tools table

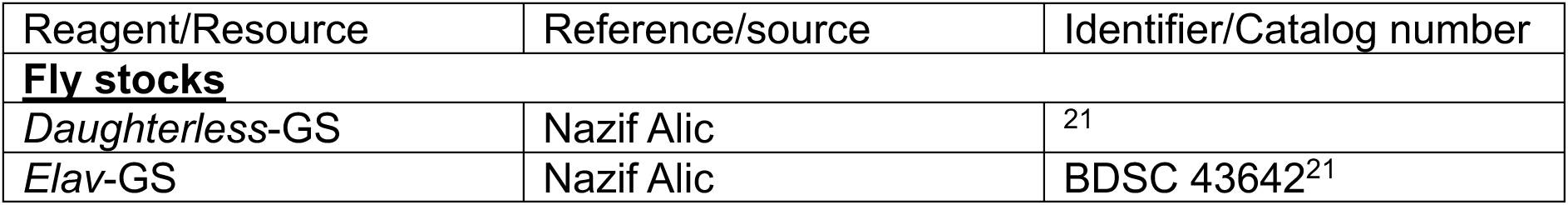
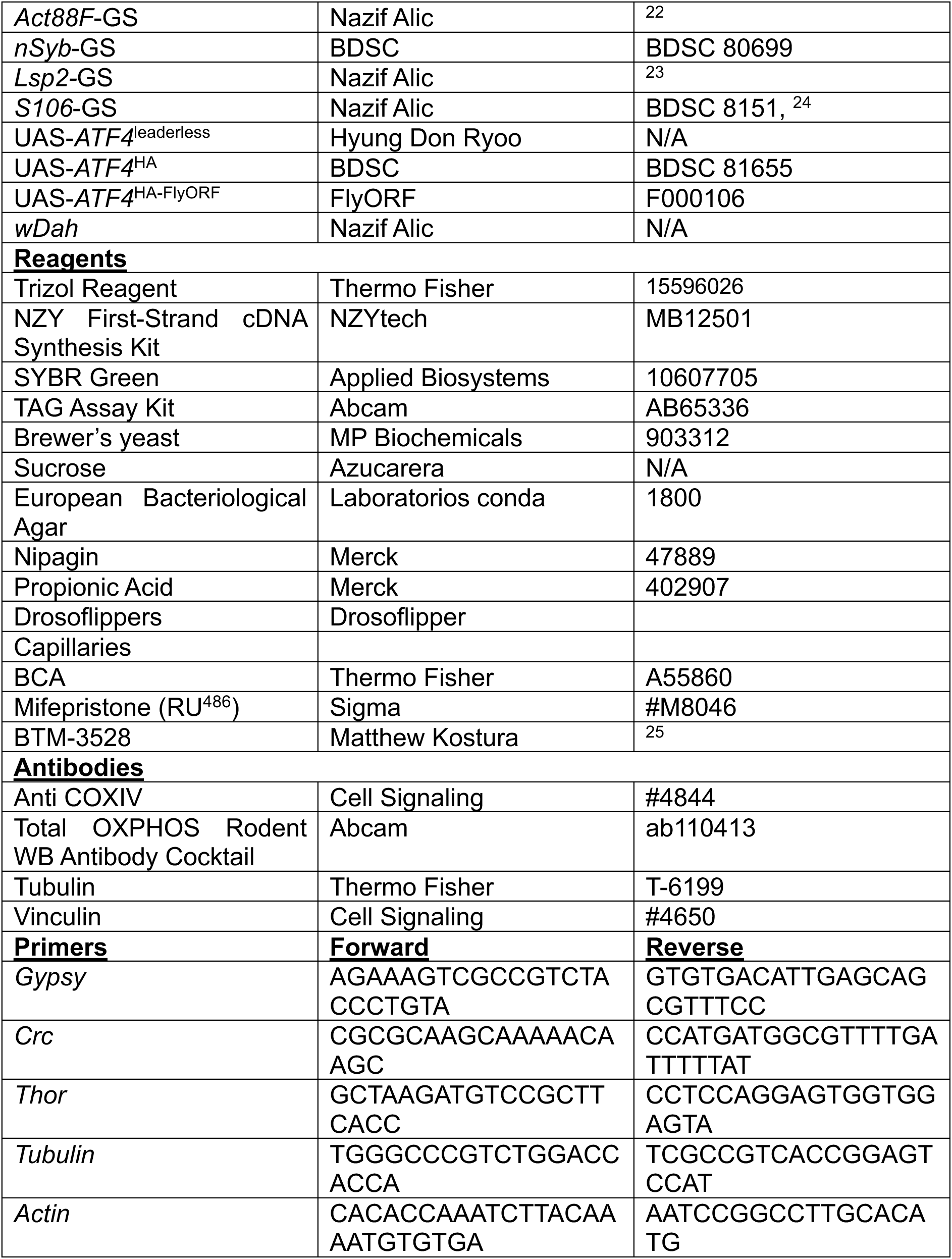

### Fly Strains and Maintenance

The *Dahomey* wild-type stock (a kind gift from Nazif Alic) was collected in 1970 in what is today Benin and has been maintained in large population cages on a 12L:12D cycle at 25°C. The *white Dahomey* (*wDah*) stock was derived by incorporation of the *w¹¹¹⁸* mutation, into the *Dahomey* background by extensive backcrossing. The *wDah*population used is *Wolbachia*-free and females from this background were used in all crosses to generate experimental flies. Mutants and transgenes were backcrossed into *wDah* for at least six generations. Unless otherwise stated, once-mated female flies were used and kept at 25°C on a 12h:12h light/dark cycle at constant 60% humidity on standard 5% sucrose (w/v), 10% brewer’s yeast (w/v) and 1.5% agar (w/v) (SYA) medium^26^. All fly stocks and transgenes used in this study are listed in the Reagents and tools table.

### Lifespan Assays

Lifespan assays were performed as described before^5^. In brief, crosses were set up in cages containing grape juice agar and live yeast, flies were allowed to mate and then embryos were collected for < 22-hr, washed in PBS and seeded into bottles at ∼22 μl per bottle to achieve standard density. Females, allowed to mate for 48h, were used in experiments, unless otherwise noted. Flies were subsequently lightly anaesthetized with CO_2_ and split into 15 per vial. Experimental vials contained either food with RU (RU+) at 200μM RU^486^ or control food (RU-, containing the equivalent volume of EtOH vehicle) according to the experiment’s design. Vials were kept in DrosoFlippers (drosoflipper.com) and flies were transferred to fresh food three times a week, when deaths/censors were recorded. Data were collected using Excel. Survival curves were compared by log-rank test. For combined analysis of independent trials, mixed-effects Cox proportional hazards models were fitted with the coxme package in R, with RU_486_ treatment as a fixed effect and experimental trial as a random intercept. Hazard ratios were estimated across all ages and within age windows (after RU_486_ withdrawal, days 40**-**60 and >60 days), with deaths beyond the end of each window treated as censored. Hazards were not proportional over age (Schoenfeld test, p < 0.05), so hazard ratios are also reported within age windows. Specificity of the response to the ATF4 transgene was tested with an RU:genotype interaction term in a model including driver-alone controls.

### Body weight measurements

At each timepoint, females were sampled at random from the experimental cohort, briefly anaesthetised with CO_2_ and weighed in groups of 3-13 females (usually groups of 5) on a precision balance METRIA Plus, Labbox; readability: 0.1 mg. Weight is expressed per fly. Up to 13 groups per condition per timepoint were weighed. Induced and uninduced flies were compared at each timepoint by two-sided Welch’s t-test.

### Capillary feeding (CAFE) assay

Food intake was measured by capillary feeder (CAFE ^27^) assay. Individual flies were housed in vials containing 1% agar for humidity and fed 5% sucrose, 5% yeast extract from calibrated 10 µl glass capillaries (71900-10, Kimble), replaced every 24 h. Consumption was corrected for evaporation using fly-free control vials. Induced and uninduced (or vehicle) flies were compared at each timepoint by Welch’s t-test on perfly values; p-values are unadjusted.

### CO2 production assays

CO_2_ production was measured in groups of 3 flies by capillary respirometry as described previously^28^. In brief, each group of 3 flies was weighed after measurement. Treatment effects were tested by ANCOVA on log(CO_2_) with log(body weight) as covariate (parallel-slopes model), fitted as a linear mixed-effects model with replicate as a random effect. Weight-adjusted means were evaluated at the mean body weight of each week.

### Fecundity assays

Egg laying was assayed at weekly intervals throughout adult life. At each timepoint, females were sampled at random from the experimental cohort, housed in vials of five on fresh SYA (RU+ or RU-, matching their lifespan treatment) and allowed to lay for 24 h. Females were then removed and the eggs per vial counted, together with the number of females present, so that fecundity was expressed as eggs per female per 24 h. Induced and uninduced flies were compared at each timepoint by two-sided Student’s t-test, pooling the two cohorts. Fecundity data are provided in Table S1, S3, S4 and S5.

### TAG quantification assays

Homogenates for TAG quantification assays were performed as shown in ^29^. Briefly, groups of 3 flies were homogenised in 100μl of 5% Triton X-100 in ddH2O, homogenised with a pestle, and heat to 95°C for 5 min. Then, they were centrifuged at 12000g for 5 min, and supernatants were collected for processing. TAG was measured with the TAG Assay Kit (Abcam, AB65336) following the manufacturers’ instructions. Values were normalised to body weight, as indicated. Starvation-induced loss was analysed with a linear model on log-transformed values including an RU:starvation interaction.

### Negative geotaxis (climbing) assays

Climbing assays were performed as described previously^5^. In brief, at the indicated timepoints flies were transferred to empty vials mounted in DrosoFlippers, giving a climbing distance of two vial heights. The same cohorts were assayed repeatedly throughout the experiment. After acclimatising for 30 min, flies were gently tapped to the bottom of the vials and video recorded for 30 s. Stills taken at a fixed timepoint (15 s, when young wild-type flies approach maximum height) were analysed in Fiji, and fly coordinates were exported to R (R Core Team). Flies whose height could not be determined were excluded, as were the top and bottom 5 % of values per condition per timepoint to limit the influence of outliers. Data were analysed with a mixed-effects linear model with flipper as a random effect, and two-sided F-test P values obtained with the *anova()* function are reported. Climbing data are provided in Table S1, S3 and S4.

### Stress assays

For oxidative stress, flies were transferred to food containing 20 mM paraquat (cat. no. 856177-1G, Merck) in SYA food. For starvation, flies were transferred to 1% agarose. Deaths were scored at least twice a day and survival was compared by log-rank test.

### RNA-seq sample preparation

Midguts, fat bodies (associated with the abdominal cuticle), thoracic muscles, and fly heads were dissected in ice-cold PBS and placed directly into ice-cold TRIzol. Three to four biological replicates were obtained per condition, each containing 12 dissected tissues. RNA was extracted in four extraction batches by Trizol-chloroform extraction, quantified on a NanoDrop 2000c spectrophotometer and eluted in nuclease-free water for subsequent quality control and sequencing (see below).

### RT-qPCR analysis

RNA was extracted with TRIzol, and cDNA was synthesised with the NZY First-Strand cDNA Synthesis Kit (NZYtech). qPCR used SYBR Green (Applied Biosystems) on a QuantStudio 7. Expression was calculated by the standard curve method, normalised to *tubulin* (*gypsy*), or to the geometric mean of *tubulin* and *actin* measured on two plates from the same cDNA (BTM organ qPCR). Primer sequences are listed in Reagents and Tools Table.

### S2R+ cell culture and mitochondrial morphology

*Drosophila* S2R+ cells were maintained at 25 °C in Schneider’s medium (Biowest #L0207) with 10% FBS (Merck # 12103C) and 1% Penicillin / Streptomycin (GIBCO, # 15070063). Cells were treated with 10 µM BTM-3528 or same volume vehicle (DMSO) for 4 h, stained with 200nM of MitoTracker Green (Thermo Fisher # M46750) at 25 °C for 30 min. Cells were washed 3 times with pre-warmed PBS and immediately visualized by live confocal microscopy using an Andor Dragonfly 505 spinning-disk system. Mitochondrial morphology was quantified using the Mitochondrial Analyzer V2.3 plugin for Fiji/ImageJ^30^. Aspect ratio, form factor and mitochondrial density were used in order to determine mitochondrial phenotype. Data are shown as SuperPlots^31^, with statistics performed on per-replicate means (n = 90 control and 96 BTM-treated images analysed in 3 independent biological replicates).

### RNA-seq of S2R+ cells

Cells were treated with 10 µM BTM-3528 or vehicle (DMSO) for 6 h (5 replicates per condition), then they were pelleted at 500 × g for 5 min, resuspended in DNA/RNA Shield (Zymo Research # R1100-50) and sent to Plasmidsaurus for RNA extraction, 3′-end library preparation and single-end sequencing. Two BTM-treated libraries yielded approximately 82,000 reads each (fewer than 10,000 deduplicated aligned reads, <0.1% of the median depth of the remaining libraries, which ranged from 5 to 16 million) and were excluded before differential testing, leaving five vehicle- and three BTM-treated libraries. Reads were processed and analysed with the same pipeline and thresholds as the fly datasets.

### mRNA library preparation and sequencing

Stranded poly(A) mRNA libraries carrying unique molecular identifiers (UMIs) were prepared and sequenced on Illumina NovaSeq instruments. The *DaGS* dataset was sequenced as paired-end 150 bp reads (NovaSeq 6000) across four flowcells with four lanes per flowcell, outsourced to the *Centro Nacional de Análisis Genómico* (CNAG); libraries carried a dual UMI encoded in the read header. The *nSyb* dataset was sequenced as single-end 100 bp, 3’-end reads (NovaSeq X Plus) in two runs, one per genotype, outsourced to Plasmidsaurus, with a single UMI encoded in the read header. For *DaGS*, reads from all lanes and flowcells were concatenated per library prior to processing. Adapter and quality trimming were performed with fastp (v0.22.0) using poly-X tail trimming, 3′ quality trimming (--cut_tail, mean quality 15) and a minimum retained read length of 50 bp; per-library QC reports were aggregated with MultiQC.

### Alignment, deduplication and counting

Trimmed reads were aligned to the *Drosophila melanogaster* BDGP6 genome with STAR (v2.7.10b) using the FlyBase r6.64 annotation and ENCODE-standard alignment parameters. A single genome index was used for both datasets to ensure identical reference handling. PCR duplicates were collapsed using the UMIs with UMI-tools dedup (v1.1.6), in paired mode for *DaGS*. Exon-level read assignment to genes was performed with featureCounts (v2.0.1); library strandedness was determined empirically by comparing assignment rates across the three strand settings, which identified forward-stranded assignment for both datasets.

### Differential expression analysis and pathway enrichment

Count matrices were analysed in R with DESeq2 (v1.50.2), using local dispersion fitting. For the *DaGS* dataset, models were fitted per organ × timepoint stratum with a fixed design (∼ rna_batch + genotype + ru + genotype:ru); both genotypes were co-sequenced, so they were pooled within a stratum and the driver-only response recovered as a linear contrast of the treatment and interaction terms. Each stratum was screened before fitting for rank deficiency, variance inflation on the treatment contrast (VIF ≤ 5) and residual degrees of freedom (≥ 6). For the *nSyb* dataset, genotype is confounded with sequencing run, so each genotype was modelled separately per tissue (∼ ext_batch + ru, reducing to ∼ ru where extraction batch was not estimable).

In both datasets, ATF4-specific genes were defined by direction-aware subtraction of the driver-only response: a gene was removed from the ATF4 list only where the control responded in the same direction and exceeded the fold-change threshold; genes responding in opposite directions were retained. Genes were retained for testing if they reached at least 10 counts in a minimum number of samples per stratum. Differential expression was called on maximum-likelihood log2 fold changes at Benjamini–Hochberg adjusted p < 0.01 and |log2 fold change| > 0.585 (1.5-fold); ashr-shrunk fold changes were used for visualisation and for GSEA ranking. One library was excluded from the *DaGS* dataset and three from the *nSyb* dataset on documented quality grounds (see Sample numbers and exclusions). Gene Ontology enrichment used topGO (v2.62.0) with a minimum node size of 10, testing genes separately against the genes tested in that stratum as background; terms were filtered at classic Fisher p < 0.05 and ordered by the weight01 algorithm, which accounts for the GO hierarchy. KEGG gene-set enrichment used clusterProfiler (v4.18.4); gseKEGG, organism "dme", gene sets of 10–500 genes) on genes ranked by ashr-shrunk log2 fold change, at FDR < 0.05. Transcription factor motif enrichment used RcisTarget with the dm6 5 kb-upstream 11-species motif rankings (mc8nr), the top 1% of the gene ranking and a normalised enrichment score threshold of 3.0; motifs were annotated to transcription factor families and summarised as the maximum NES per family.

### Sample numbers and exclusions

Principal component analysis was performed on variance-stabilised counts using the 500 most variable genes, separately for each stratum × genotype combination rather than on pooled data. All sequenced libraries are shown, including those subsequently excluded from differential testing. Of 128 dissected samples in the *DaGS* dataset, 117 yielded RNA suitable for library preparation and sequencing. Losses were concentrated in one extraction batch but were balanced across genotype and treatment and therefore did not bias treatment contrasts. One muscle library was excluded as a technical outlier, showing markedly reduced sample**-**sample network connectivity within its own genotype (Zk = −2.43, against −0.06 to +0.49 for the other seven **same-stratum** libraries; permutation p = 0.002) with no evidence of a treatment-label problem. Its removal changed differential expression counts for that stratum within the range spanned by leave-one-out resampling (309–543), so the exclusion is both well evidenced and low in consequence. Four further libraries were assessed and retained, none having independent technical support for removal. One stratum, fat body at the post timepoint, is underpowered: leave-one-out differential expression counts range from 35 to 1776 against 42 with all samples, compared with 1.8–3.2-fold variation in every other stratum. It is reported with all samples retained and this sensitivity range disclosed, rather than stabilised by exclusion.

For the *nSyb* dataset, 64 libraries were prepared (four biological replicates per stratum) and three were excluded. One fat body library (ATF4, uninduced) was excluded following an annotation error identified in the dissection records. One head library (ATF4, uninduced) was excluded because it yielded approximately half the sequencing depth of the other head libraries (4.1 versus 8.4–10.3 million reads), with fewer detected genes, and clustered with the induced group in principal component analysis. One muscle library (driver alone, induced) was the only sample of the 64 flagged by a permutation-calibrated connectivity test (p < 0.001), with a 4.6-fold reduction in dispersion on its removal (0.1646 to 0.0357). Four further libraries flagged in an earlier audit on principal component position alone were reinstated after re-assessment against permutation-calibrated connectivity and dispersion criteria, none showing a measurable effect on dispersion. All exclusions were made before differential testing. Following these removals, the fat body and head ATF4 models and the muscle *w*Dah model retain two residual degrees of freedom, so dispersion estimates for those strata are correspondingly less well determined. Uninduced fat body libraries contained slightly more gut-derived transcripts than induced ones. Excluding gut-enriched genes reduced the number of differentially expressed genes but did not change the overall pattern of the response, and transposable element results were unaffected.

### Transcription factor binding site enrichment analysis

Putative transcriptional regulators of ATF4 target genes were identified using RcisTarget (v1.20.0) with the *Drosophila melanogaster* dm6 motif rankings database (5 kb upstream of transcription start sites, 11-species conservation; dm6-5kb-upstream-full-tx-11species.mc8nr). For each tissue and timepoint combination, ATF4-specific upregulated and downregulated gene lists were analysed separately; gene sets with fewer than 10 genes present in the database were not tested. Motif enrichment was assessed at a normalised enrichment score (NES) threshold of 3.0, with the AUC calculated over the top 1% of database gene rankings. Enriched motifs were assigned to transcription factor families by ordered regular-expression matching on the transcription factor name parsed from the motif identifier, after removal of species and variant suffixes. For each family, enrichment was summarised per tissue, timepoint and direction as the maximum NES across all motifs annotated to that family, so that a family is scored by its most strongly enriched motif irrespective of how many motifs it has in the database, and displayed as a heatmap.

### Transposable element quantification and abundance analysis

Transposable elements are absent from the FlyBase r6.64 gene annotation, so the gene-level count matrices contain no TE signal and TE abundance was quantified separately. A combined index of the r6.64 transcriptome and the r6.64 transposon insertion set was built with salmon (v2.6.0, k = 31), so that TE-derived and gene-derived reads compete for assignment; indexing TEs alone causes gene-derived reads resembling TE sequence to be force-assigned to TEs and inflates apparent abundance. Reads were quantified with automatic library-type inference, selective alignment, and sequence-specific and GC bias correction. The FlyBase transposon reference describes 5,734 individual insertions across 125 families; because copies within a family are near-identical and 2,052 insertions are shorter than 200 bp, per-insertion estimates are not identifiable from reads of this length, and insertion-level estimates were summed to family level. Gene-level counts from the same quantification were used to derive size factors.

TE abundance was analysed in fat body, separately in each genotype arm. As the primary test, total TE burden was summarised as one value per pool (summed TE counts divided by summed gene counts) and compared between treatment groups within each genotype by Welch’s t-test, with a Wilcoxon rank-sum test reported alongside. Because normality cannot be assessed at this sample size and the rank test has an arithmetic floor at this n, both are reported together with that floor.

As a confirmatory test, per-family fold changes were estimated with DESeq2 and the median log2 fold change across families compared against a null generated by reassigning treatment labels between pools. All possible label assignments were enumerated exhaustively rather than sampled, giving an exact p-value; the number of available arrangements bounds the minimum attainable p-value, which is reported as such where reached. Because each permuted fit preserves the family-to-family correlation structure of the observed data, no assumption of independence between families is required. Size factors were estimated from gene counts and applied to the TE matrix, since median-of-ratios normalisation on TE counts would assume most families are unchanged. Dispersions were estimated once and held fixed across permutations, with the same family set tested throughout. Families were retained if they reached at least 10 counts in as many pools as the smaller treatment group.

The number of families changing in each direction is reported descriptively only; no test treating families as independent replicates was applied, since such a test uses the number of annotated families rather than the number of animals as its sample size. The *wDah* control was tested in parallel rather than subtracted, as subtraction would add its variance to the ATF4 estimate. The two genotypes were sequenced in separate runs and sit at different absolute TE fractions, so only within-genotype changes are interpretable.

### Respirometry in frozen fat bodies

Respiratory chain capacity was measured in frozen fat body homogenates as described elsewhere^32^. Abdominal fat bodies from 12 females per cohort were dissected, snap-frozen and homogenised in 300 µL MAS buffer (70 mM sucrose, 220 mM mannitol, 5 mM KH_2_PO₄, 5 mM MgCl_2_, 1 mM EGTA, 2 mM HEPES, pH 7.2). To permeabilise mitochondrial membranes and allow substrate access, homogenates underwent three freeze-thaw cycles (between liquid nitrogen and 37 °C) prior to assaying. Additionally, alamethicin (Fisher Scientific, AAJ63829MCR) was added to the MAS buffer on the day of the assay. 2 µg protein were loaded per well of an XFe96 plate (Agilent). Complex I-driven respiration was measured with NADH (10.77 mM), and complex II-driven respiration with succinate (5 mM) plus rotenone (0.2 mM), in the presence of cytochrome c (10 µg/ml). Sequential injections of antimycin A (4 µM), TMPD (0.5 mM)/ascorbate (1 mM) and azide (50 mM) were used. Complex I or II capacity was calculated as substrate minus antimycin A, and complex IV as TMPD/ascorbate minus azide, normalised to protein. Wells in which the second substrate measurement fell below 70% of the first (substrate depletion; 6 wells, all succinate) were excluded from complex II quantification. Technical wells were averaged within each cohort, and cohorts were compared by Welch’s t-test (n = 4).

### BTM feeding

BTM-3528 was added to SYA at 10 µM from a 30mM DMSO stock, with vehicle-matched control food. Treatment windows matched those of the genetic inductions (1 week, 3 weeks and chronic). For target engagement, organs were dissected after 24 h of exposure and analysed by qPCR.

### Western-blotting

Western blots were performed in the same homogenates used for respirometry. 20µg of protein, quantified by BCA following manufacturer’s instructions, were resolved by gradient SDS-PAGE and transferred to a PVDF membrane using manufacturer’s instructions. Membranes were probed with antibodies against COX4 (1/1000) and Total OXPHOS Rodent WB Antibody Cocktail (1/1000, which detected ATP5A only, see Reagents and tools table). Secondary antibodies were conjugated with fluorescent probes (LiCOR) and blots were developed in an OdysseyXF Imaging System. Signals were quantified in Fiji and normalised to vinculin or tubulin. Raw western blot images are shown in Table S6. Groups were compared with Welch’s t-test.

### Data availability

RNA-seq data have been deposited in the Gene Expression Omnibus under accession GSE350078. Analysis scripts are available on request.

## Results

### Transient ATF4 induction in early adulthood has long lasting consequences that extend lifespan

We established a new *Drosophila* model that enabled an inducible and reversible gain-of-function of the transcription factor ATF4 (*crc* in *Drosophila*) *in vivo* (Figure 1A). These lines consist of a previously validated line lacking the leader sequence (UAS-*crc*^leaderless^), namely the 5′ upstream open reading frames that constitutively repress ATF4 translation. Therefore, this line shows an increase in ATF4 function independently of eIF2α phosphorylation^33,34^. Validating our success increasing ATF4 function in this line, expression of *crc*^leaderless^ significantly upregulated the transcription of the canonical ATF4 target *thor*/4E-BP (Figure S1A).

**Figure 1.**
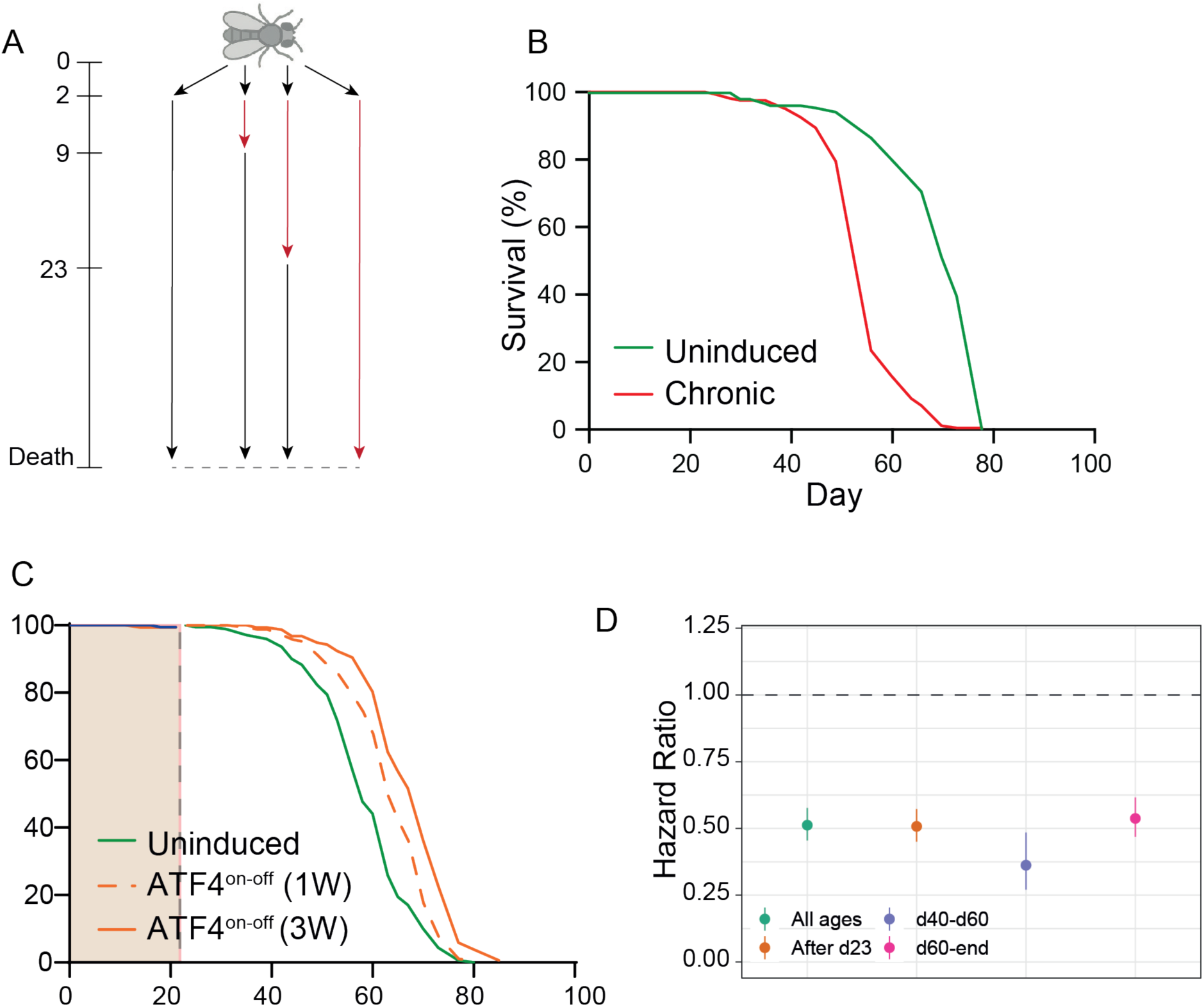
**a)** Schematic diagram of the experimental setup. The period of RU feeding is marked in red, and food without RU, in black. ATF4^on-off^ flies are induced until day 23 and then transferred to control food thereafter. **b)** Adult-only, chronic ATF4 overexpression lifespan. Uninduced n = 155 dead/3 censored, chronic n = 158 dead/6 censored, p = 2.3 x 10^-39^, log-rank test. Driver-alone control is shown in Figure S1I. **c)** ATF4^on-off^ and ATF4^on-off^ (one week) lifespan. Uninduced n = 170 dead/8 censored, ATF4^on-off-1W^ n = 165 dead/5 censored, ATF4^on-off^ n = 156 dead/17 censored, p = 4.23 x 10^-5^ (ATF4 ^on-off-1W^ vs uninduced) 3.63 x 10^-15^ (ATF4^on-off^ vs uninduced) and , log-rank test. **d)** Hazard ratios (points) and 95% CIs (whiskers) for ATF4^on-off^ vs uninduced flies, from mixed-effects Cox proportional hazards models with experimental trial as a random effect, fitted on the combined data of four independent trials (1166 deaths, 114 censored). Hazard ratios were estimated across all ages and within age windows (after day 23, i.e. after RU withdrawal; days 40-60; >60 days). HRs < 1 indicate lower risk of death in ATF4^on-off^ flies. Additional individual trials are shown in S1B-D. RU_486_ had no effect in driver-alone controls (Table S1).

Then, we employed the GeneSwitch (GS) system^35,36^, to reversibly increase ATF4 function in a temporary manner in the whole organism, with the driver *Daughterless-*GeneSwitch (*Da*GS). We named this new model ATF4^on-off^. Restricting ATF4 induction to adulthood avoided the developmental lethality caused by ATF4 induction in the larval stage^37^. Consistent with recent evidence^38^, we reproduced using our line that chronic ubiquitous ATF4 activation throughout adulthood shortened lifespan (Figure 1B). To test whether the length of induction determines its outcome, we restricted ubiquitous and transient ATF4 activation to early adulthood, with the activation lasting either one or three weeks. Both transient activation paradigms extended lifespan, with the 3-week activation being more efficacious than 1-week (Figure 1C). Importantly, this lifespan extension was consistently confirmed across three independent biological replicates of the 3-week induction (Figure S1B-D). Transient activation restricted to midlife also extended lifespan, although less markedly than in early adulthood (Figure S1F), indicating that the window of sensitivity is not confined to the first weeks of adult life. Importantly, two additional and independent UAS-ATF4 lines^39^ reproduced the lifespan extension (Figure S1G and H). Notably, lifespan extension was restricted to females (Figure S1E). We therefore used the three-week induction in females for all subsequent ubiquitous experiments, which provides the greater dynamic range to investigate the mechanisms responsible. We combined data from four independent trials (1166 deaths) in a mixed-effects Cox proportional hazards (CPH) model. ATF4^on-off^ approximately halved the risk of death (Table S1), an effect that persisted into old age (Figure 1D). Altogether, these data show that increasing ATF4 activity confined to specific windows of early adulthood is sufficient to promote longevity, and that the length of its gain-of-function is a key determinant of its actions on lifespan.

### Transient whole body ATF4 induction improves healthspan, associated with decreased body protein content and egg-laying

We next aimed to determine whether life-span extension induced by ATF4^on-off^ was concurrent to improved health outcomes. Thus, we next studied two key phenotypic traits of healthy ageing: body composition and neuromuscular activity. We found that ATF4^on-off^ flies were significantly lighter during the induction or ATF4^on^ period, but this difference in body weight normalised soon after ATF4 induction was switched off (Figure 2A). This weight loss could not be explained by reduced feeding, as food intake was unchanged or even slightly increased during induction (Figure S2A).

**Figure 2.**
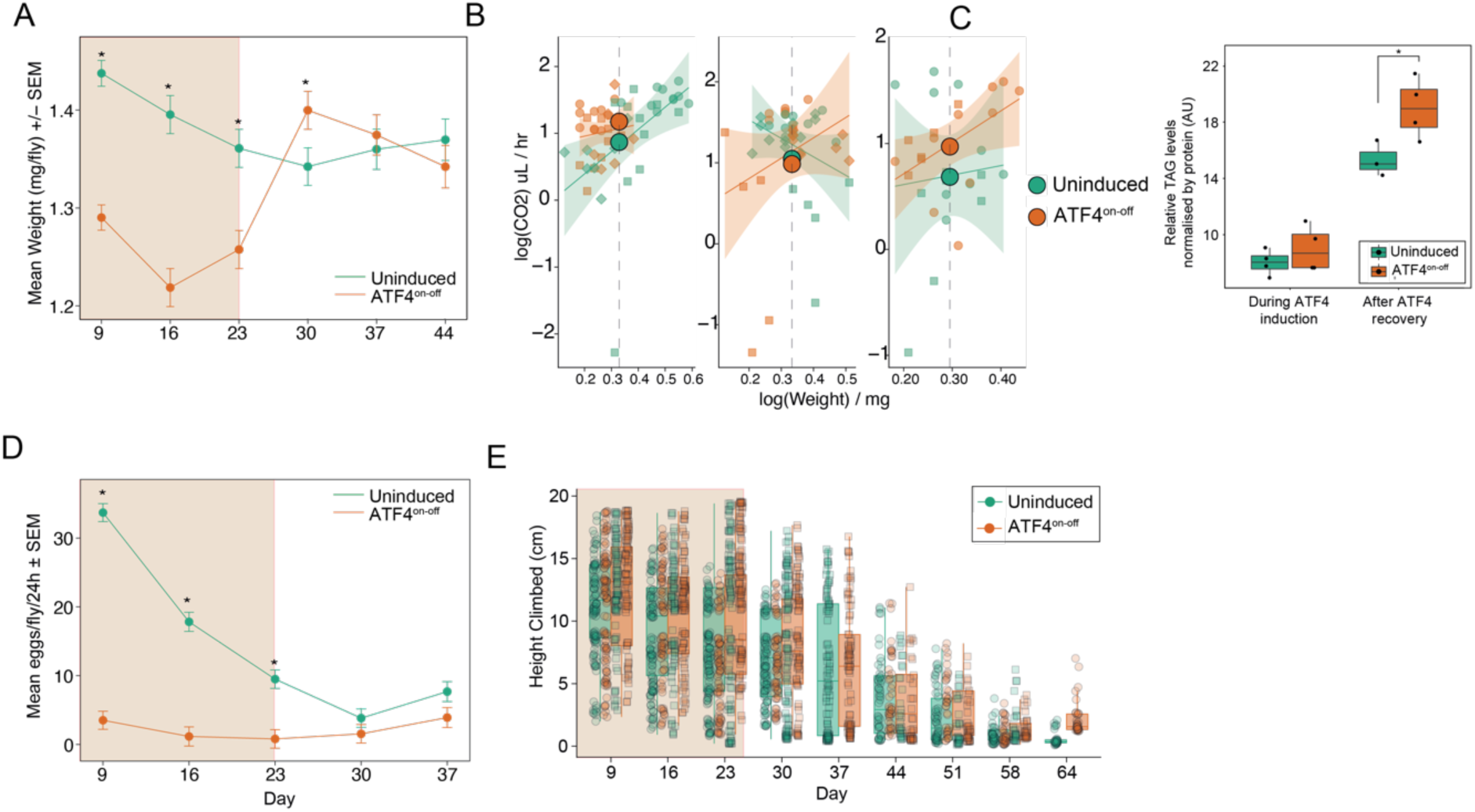
a) Body weight of whole ATF4on-off and uninduced flies throughout life. Mean ± SEM. * p < 0.05, Welch’s t-test per timepoint. n = 5-21 flies per condition per timepoint, 2 biological replicates. b) CO_2_ production per fly at weeks 1, 4 and 6. Points represent groups of 3 flies; lines show fitted allometric relationships between log(CO_2_) and log(body weight) per treatment group, with 95% confidence intervals (shaded). Large filled points indicate weight-adjusted means from ANCOVA (parallel slopes), evaluated at the mean body weight of each week. No significant treatment effect at week 1 (p = 0.132), week 4 (p = 0.394) or week 6 (p = 0.192); linear mixed-effects models with replicate as a random effect (3 biological replicates, n = 15-28 flies per condition per week). c) TAG/protein ratio during ATF4 induction (week 1) and after recovery (week 5). Box plots show median and quartiles; points represent individual samples (n = 3-5 per condition). Week 1: p = 0.35; week 5: p = 0.038, t-test. d) Fecundity of ATF4on-off and uninduced flies, expressed as eggs per female per 24 h. Mean ± SEM. * p < 0.05, Welch’s t-test per timepoint. n = 4-5 vials per condition, 2 biological replicates. e) Height climbed by ATF4on-off and uninduced flies, from 2 independent trials. n = 88-108 flies per condition. Box plots: quartiles; whiskers: extremes; overlay: individual flies. Effect of ATF4^on-off^ p = 0.0006, age p < 0.0001, age:ATF4on-off interaction p = 0.134, mixed-effects linear model. In a, d and e, the shaded area represents the period of RU486 exposure.

To determine whether the initial weight reduction was caused by an increase energy expenditure, we subsequently measured life-course CO_2_ production rates *in vivo* ^28^ in ATF4^on-off^ flies at 3 different time points: during direct ATF4 induction (week 1), one week after ATF4 was off (week 4), and at old age (week 6). Overall, we did not observe any increase in CO_2_ production rates when modelled at the same body mass between the two groups using ANCOVA (Figure 2B). This weight loss during induction is therefore not explained by increased whole body oxidative metabolism to CO2. As we measured CO2 production, but could not measure O2 consumption in intact flies, there was still the possibility that energy expenditure was elevated by an increase in fat oxidation.

To overcome this limitation and explore the basis of this transient weight reduction, we determined their body composition. This was achieved by measuring total triacylglycerol (TAG) and protein content in whole fly lysates, during and after ATF4 induction. During induction (ATF4^on^), total TAG and the TAG/protein ratio were unchanged, but total protein was reduced (Figure 2C, S2B). After 12 days of ATF4^off^, body weight had returned to control levels. However, these ATF4^off^ flies showed a higher TAG/protein ratio (Figure 2C), driven by lower protein content (Figure S2B) rather than increased TAG (Figure S2C). This shift in body composition, which emerged only after ATF4 had been switched off, indicates that transient ATF4 activation leaves a persistent metabolic imprint consistent with reduced protein synthesis, rather than increased fatty acid oxidation.

Given that egg-laying is an energy consuming process previously shown to shorten lifespan, and that we found that ATF4 ^on-off^ improved lifespan exclusively in females, we determined whether ATF4^on-off^ decreased egg-laying. Consistent with this hypothesis, we observed a dramatic reduction of egg-laying in ATF4^on-off^ flies compared to controls during the *on* period (Figure 2D). Accordingly, the ISR has recently been shown to regulate fecundity non-autonomously, by acting in the fat body through ATF4^40^. However, when we activated ATF4 exclusively in this the fat body with *Lsp2-*GS, egg laying was unchanged (Figure S2D). The reduction in egg laying is therefore likely to arise from ATF4 activity in another tissue. Strikingly, egg-laying partially caught up to the controls when the ATF4 *off* period was set, showing that reduction in egg-laying is in itself transient.

We next studied ATF4^on-off^ flies’ neuromuscular health by climbing assays, to establish whether the decrease in total protein content was associated with altered muscle function, a major protein reservoir^41^. ATF4^on-off^ flies climbed significantly better than controls, both during induction and after ATF4 was switched off, and this advantage was maintained throughout life (Figure 2E). Of note, no effects on lifespan, body weight, food intake or climbing were observed in driver-alone or UAS-*ATF4*-alone controls (Figure S2E-J), confirming that the phenotypes of ATF4^on-off^ flies result from transient ATF4 induction. Taken together, these results show that transient ubiquitous ATF4 activation in early adulthood improves lifespan and locomotor health, while inducing persistent changes in body composition.

### Transient whole body ATF4 activation induces lasting transcriptional programs of redox, lipid, protein and oxidative metabolism in a tissue-specific manner

Because transient activation of a transcription factor can establish lasting transcriptional states^5^, we analysed the transcriptome of four major organs that determine health span in aged flies: we performed RNA-seq of the midgut, fat body, thoracic muscle and the head. These analyses were performed during ATF4 induction (7 days of ATF4^on^) and 12 days after ATF4 was switched off (Figure 3A). We confirmed the upregulation of ATF4 expression (*crc*) and its previously reported downstream targets *thor* and *scyl*^19,42,43^ during induction in these organs, and their return to basal levels after ATF4 was switched off (Figure S3A). To differentiate genes specifically changed by ATF4 activity, versus genes changed by RU_486_ treatment, we removed genes that responded in the same direction in driver-alone controls treated with RU_486_ (Table S2; see Methods).

**Figure 3.**
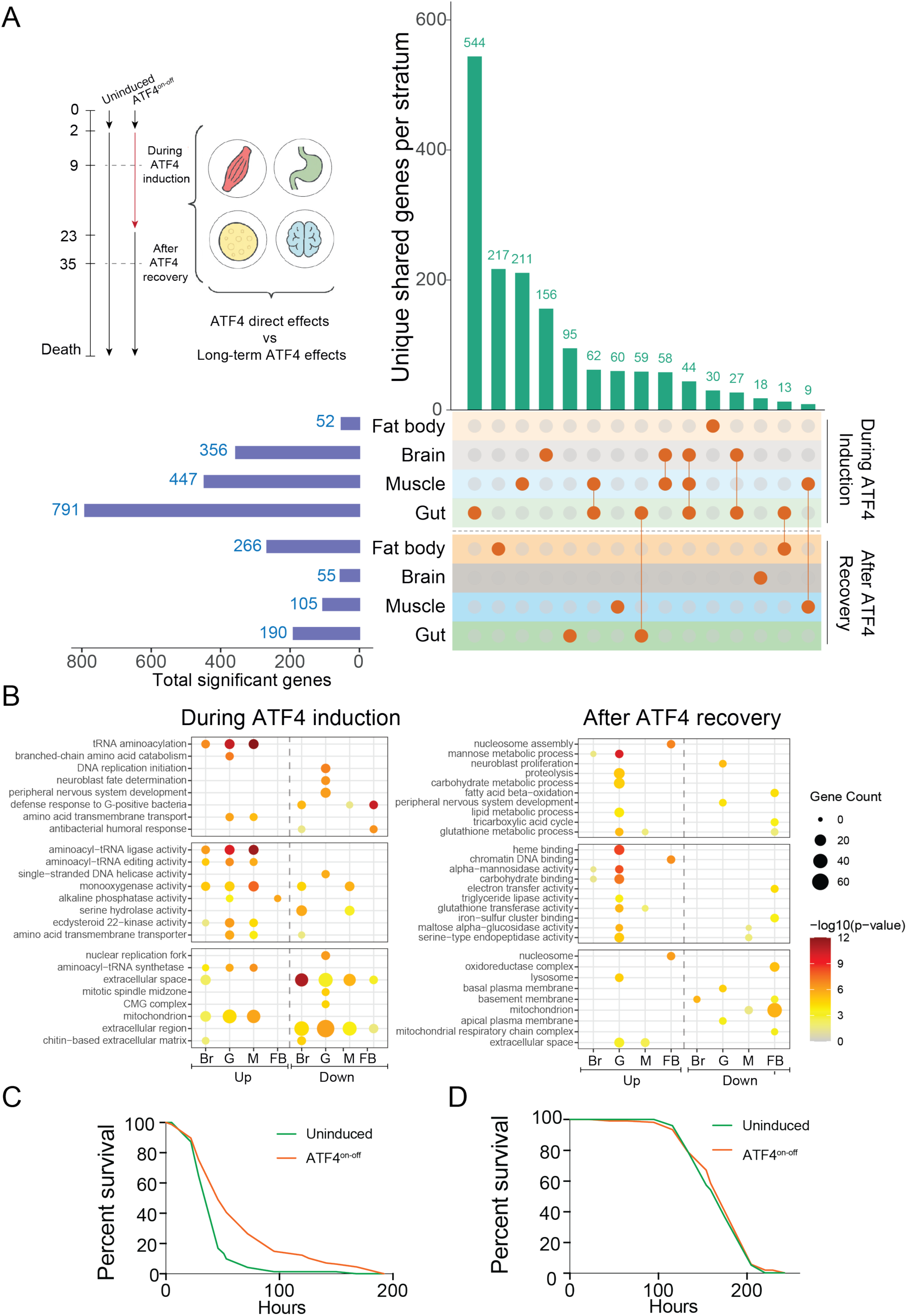
**a)** Experimental setup of the tissue and time-specific conditions analysed by RNA-seq. The same cohort of flies were dissected on day 9 (during ATF4 induction), and on day 35 (after ATF4 recovery, 12 days after RU withdrawal), and Upset plot showing the overlapping genes between datasets (green bars), with the total number of genes per condition (blue bars). Only the interactions with 9 or more genes are shown. **b)** Top 8 overrepresented GO terms per condition, both in the upregulated (Up) and downregulated (Down) genes. H: Head, G: gut, M: muscle and FB: fat body. The complete list of significant GO terms is shown in TableS2. **c)** Survival assay against oxidative stress (20mM paraquat), uninduced n = 142 dead/0 censored, ATF4^on-off^ n = 148 dead/7 censored. P = 1.005 x 10^-8^, log-rank test. **d)** Starvation survival assay (1% agarose) uninduced n = 98 dead/2 censored, ATF4^on-off^ n = 106 dead/0 censored. P = 0.48, log-rank test. Driver-alone starvation assay is shown in Figure S4F.

Using a stringent threshold (FDR < 0.01 and ≥ 1.5-fold change), we identified extensive gene expression changes during induction, which were largely reduced after recovery (Figure 3A, S3B; Table S2). Principal Component Analysis (PCA) clearly showed a separation between uninduced and ATF4^on^ samples, with this separation being less pronounced in ATF4^off^ (Figure S3B). The gene expression changes during induction (ATF4^on^), when compared to ATF4^off^, were the greatest in head, gut and muscle. The fat body showed the opposite pattern: gene expression changes were bigger in ATF4^off^ flies, when compared to ATF4^on^ flies (induction). During induction, differentially expressed genes overlapped significantly between all pairs of tissues, nearly always changing in the same direction, and 60 genes were shared by at least three tissues, 59 of them changing in the same direction (Table S2). This indicates that a shared core of ATF4 targets accompanies the tissue-specific responses. After recovery, overlap between tissues was sparse, with only one gene shared by three or more tissues (Table S2), consistent with a tissue-specific secondary response. Between timepoints, overlap was small but significant in all tissues except the fat body (Figure S4A), indicating that little of the acute response to ATF4 induction carries over once ATF4 is switched off.

Gene Ontology analysis revealed tissue-specific processes during induction and recovery (Figure 3B, Table S2). Upregulated genes in ATF4^on^ flies were enriched for glutathione metabolism, cellular oxidant detoxification and tRNA aminoacylation across tissues, consistent with the canonical ATF4 downstream transcriptional programme^6,7^, whereas immune-related genes were downregulated in fat body and head. With the exception of detoxification genes in the gut, these ATF4 induction signatures were not maintained after recovery as expected. Instead, a distinct set of gene expression changes emerged in a tissue-specific manner. Most tissues, particularly fat body and muscle, showed downregulation of metabolic pathways, including amino acid metabolism, oxidative phosphorylation and mitochondrial respiratory chain assembly. The head showed an increase in methionine metabolism and a decrease in long-chain fatty acid and triglyceride metabolism gene expression. The gut showed an opposite behaviour, with increased lipid metabolism genes, as well as showing elevated glutathione metabolism (Table S2). KEGG gene set enrichment analysis confirmed these patterns, including reduced oxidative phosphorylation in fat body and muscle after recovery (Figure S4B). Together, these data indicate a two-phase response: an acute phase of canonical stress adaptation, followed by a distinct and persistent phase of metabolic remodelling that outlasts ATF4 expression.

To identify the transcriptional regulators upstream of the genes characterising this second phase, persistent response, we performed transcription factor motif enrichment analyses. (Figure S4C). During induction, ATF4 motifs were enriched among upregulated genes, and not in downregulated genes in all tissues, except the fat body. These data confirm the validity of our motif analysis, showing successful ATF4 motif detection during ATF4 induction. During induction, tissue-specific enrichment of HSF motifs in head, gut and muscle, and of C/EBP motifs in muscle and gut, was also detected, suggesting that ATF4 engages different co-regulators depending on the tissue. After recovery, the motif landscape shifted markedly: ATF4 motifs were no longer enriched in any tissue, whereas ETS motifs were enriched in all four tissues, SOX motifs in head, gut and muscle, and GATA motifs in head and gut. This transition suggests that transient ATF4 activation initiates a transcriptional relay, maintained by other transcription factors after the initiating signal has ended.

We next asked whether this second persistent phase of changes in gene expression, associated with SOX, GATA and ETS, could provide resistance to different stressors. The acute antioxidant signature was largely lost after ATF4 was switched off, with detoxification genes remaining elevated only in the gut. Therefore, we exposed ATF4^off^ flies (12 days after switch off) to paraquat, a known oxidative agent that increases ROS and generates superoxide at complex I. ATF4^on-off^ flies were more resistant to paraquat, surviving significantly longer than controls (Figure 3C, S4D). RU alone had a smaller and inconsistent effect in driver-alone controls (Figure S4E; Table S2), and the protection in ATF4^on-off^ flies significantly exceeded it (RU:genotype interaction p = 0.003, Cox model stratified by trial). In contrast to paraquat resistance, ATF4^on-off^ flies were not more protected from the deleterious effects of starvation (Figure 3D).

Altogether, these results show that transient ATF4 activation induces an acute, canonical stress response that is replaced by a distinct, tissue-specific programme of metabolic remodelling, accompanied by lasting resistance to oxidative stress.

### One week of neuronal ATF4 activation is sufficient to extend lifespan without metabolic trade-offs

To identify the main processes contributing to lifespan extension by transient ATF4 induction early in life, we restricted ATF4^on-off^ induction to individual tissues. Because ATF4 activity is persistently increased in the livers of long-lived mice ^11^, we first targeted the fat body, the fly equivalent of mammalian liver and adipose tissue. However, neither transient nor chronic ATF4 activation in the fat body and gut (*S106*-GS) extended lifespan (Figure S5A, B), consistent with a recent report using the same driver^38^. Transient ATF4 activation in thoracic muscle (*Act88F*-GS) also failed to extend lifespan (Figure S5C, D). Three weeks of ATF4 activation in neurons (*Elav*-GS) likewise had no effect (Figure S5E). Restricting neuronal ATF4 activation to the first week of adulthood extended lifespan, reproducing the effect of ubiquitous activation with two independent pan-neuronal drivers, *nSyb*-GS^44^ (Figure 4A, S5F and G) and *Elav*-GS (Figure S5E).

**Figure 4.**
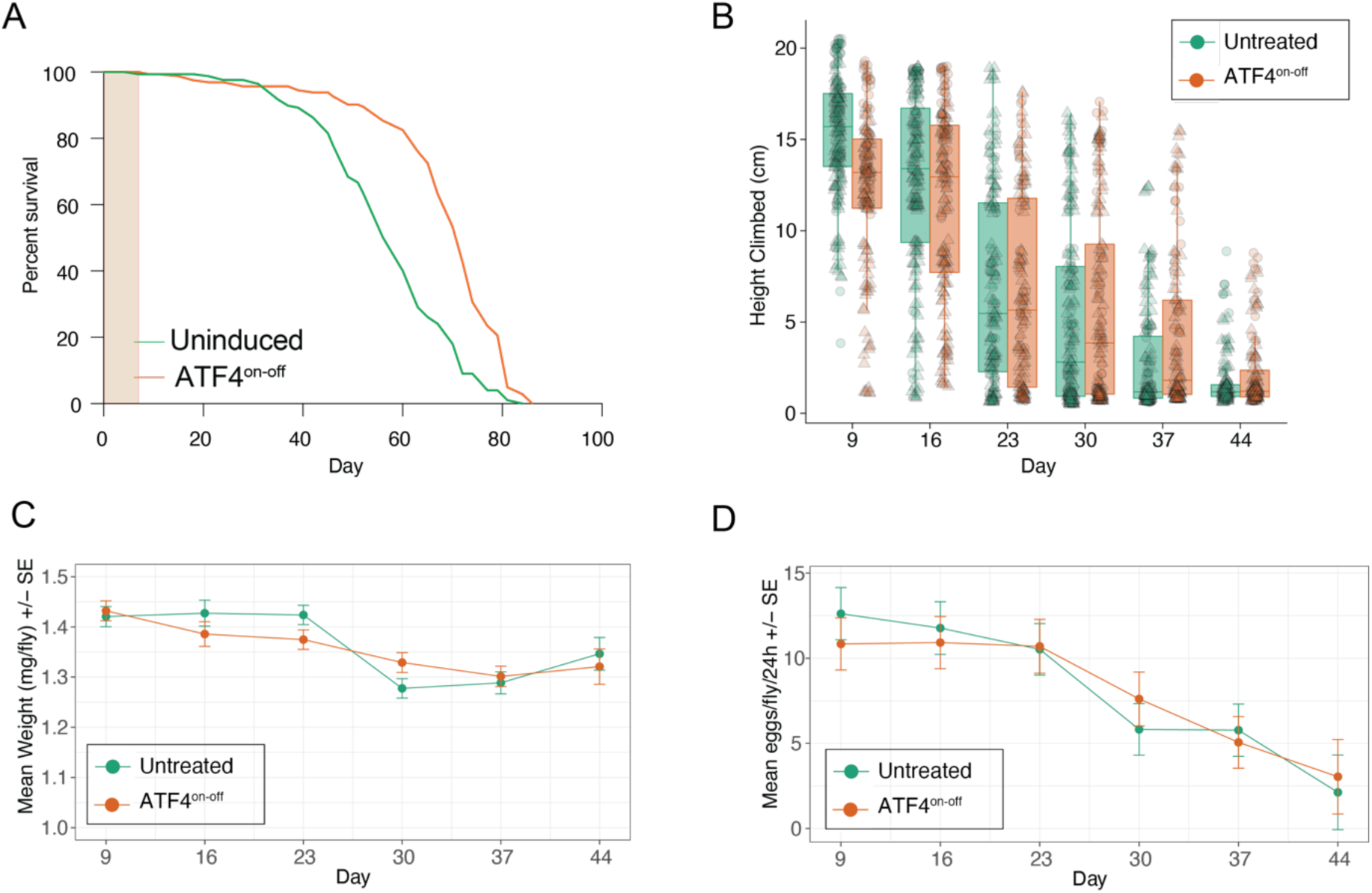
**a)** Neuron-specific ATF4^on-off^ lifespan. Uninduced n = 125 dead/43 censored, ATF4^on-off^ n = 143 dead/21 censored, p = 4.015 x 10^-14^, log-rank test. **b)** Height climbed by neuronal ATF4^on-off^ flies vs uninduced controls, from 2 independent trials. n = 63-143 per experimental condition. Boxplots: quartiles; whiskers: extremes; overlay: individual data points. Effect of RU486 p < 0.0001, age p < 0.0001, RU:age p < 0.0001; RU486:age:genotype vs driver-alone controls p < 0.0001; mixed-effects LM. **c)** Body weight of neuronal ATF4^on-off^ flies (*nSyb*-GS > UAS-*ATF4*) and uninduced controls across life. Mean ± SEM. n = 8-19 vials of 2-6 females per condition per timepoint, 2 biological replicates (day 44, one replicate). Welch’s t-test per timepoint: day 23, p = 0.029; all other days p > 0.09. **d)** Fecundity of neuronal ATF4^on-off^ flies and uninduced controls across life, expressed as eggs per female per 24 h. Points show model-estimated means ± SE. n = 10 vials of 4-5 females per condition per timepoint (day 44, 5 vials from one replicate), 2 biological replicates, Welch’s t-test per timepoint, all p > 0.26.

Beyond lifespan, one-week, neuronal ATF4^on-off^ improved age-related locomotor decline: climbing ability declined more slowly compared with the controls, with ATF4^on-off^ outperforming them in late-life (Figure 4B). In marked contrast to ubiquitous ATF4^on-off^, neuronal-restricted ATF4^on-off^ had no sustained effect on body weight (Figure 4C) or egg laying (Figure 4D). These latter data demonstrated that the decreased egg-laying and the changes in body composition induced by ATF4 induction can be uncoupled to its function in longevity. All the metabolic and health phenotypes were reproduced with a second, independent pan-neuronal driver, *Elav*-GS: one week of neuronal ATF4^on-off^ extended lifespan (Figure S5E), improved climbing (Figure S5H), and did not alter egg laying or body weight (Figure S5I, J). RU486 reduced mortality in neuronal ATF4^on-off^ flies in each of three independent trials (pooled HR = 0.55). RU486 alone altered survival of *nSyb*-GS driver-alone controls in two of the three trials, but in opposite directions (Table S3; pooled HR = 1.03), and its effect was smaller than in ATF4^on-off^ flies overall (mixed effects CPH model RU:genotype interaction, p = 1.2 x 10^-6^). With the second pan-neuronal driver, Elav-GS, ATF4^on-off^ extended lifespan whereas RU_486_ had no effect in driver-alone controls (Figure S5E, L). Similarly, the slower age-related climbing decline was specific to ATF4^on-off^ flies (RU486:age:genotype interaction, p = 0.0135; Figure S5H, N).

Together, these findings show that a single week of neuronal ATF4 activation in early adulthood is sufficient to extend lifespan and slow locomotor decline without the costs to body weight and fecundity of whole-body activation. This suggests that the changes in body composition caused by ubiquitous ATF4^on-off^ do not contribute to its benefits, and may even limit them.

### BTM-3528 activates the mitochondrial integrated stress response in *Drosophila* cells, but does not extend lifespan

Having established that a brief ATF4 pulse can install multiple durable pro-longevity programmes, we asked whether the ATF4 response could be engaged pharmacologically. To do so, we used BTM-3528 (hereafter BTM), a pyrazolo-thiazole compound that increases ATF4 activity engaging the mitochondrial OMA1-DELE-HRI axis. Accordingly, the phenotype observed in BTM-treated cells is mitochondrial fragmentation as a result of OMA1 activation^25,45^. As the orthologues of OMA1 and DELE1 in *Drosophila* are unclear and uncharacterized respectively, we first determined whether BTM treatment could induce mitochondrial fragmentation and activate ATF4 in *Drosophila* cells.

To this end, we treated *Drosophila* Schneider’s 2R+ (S2R+) cells with 10 µM BTM for 4 h, a concentration that promotes OMA1 activation-dependent fission in human cells^25^. We observed that BTM treatment induced mitochondrial fragmentation, represented by a decreased aspect ratio, form factor and increased mitochondrial density, indicating a more punctate and less interconnected network (Figure 5A and B), showing that BTM is active in fly mitochondria. Six hours of BTM treatment strongly induces ATF4 activity in human cells^25^. S2R+ cells treated for 6 hours showed extensive transcriptional remodelling, changing the expression of 1420 genes (730 up, 690 down) (Figure 5C). Upregulated genes were enriched for protein refolding and tRNA aminoacylation, while downregulated genes were enriched for DNA replication, mitosis and cell division (Figure 5D). Gene set enrichment analysis confirmed these patterns, with upregulation of aminoacyl-tRNA biosynthesis, glutathione metabolism and the KEGG ISR pathway, and downregulation of DNA replication and repair pathways (Table S5). Motif enrichment identified HSF and ATF4 binding sites among upregulated genes and E2F sites among downregulated genes (Figure 5E), and the BTM response overlapped significantly with the response to ubiquitous ATF4 induction in vivo in head, gut and muscle (Figure S6A).

**Figure 5.**
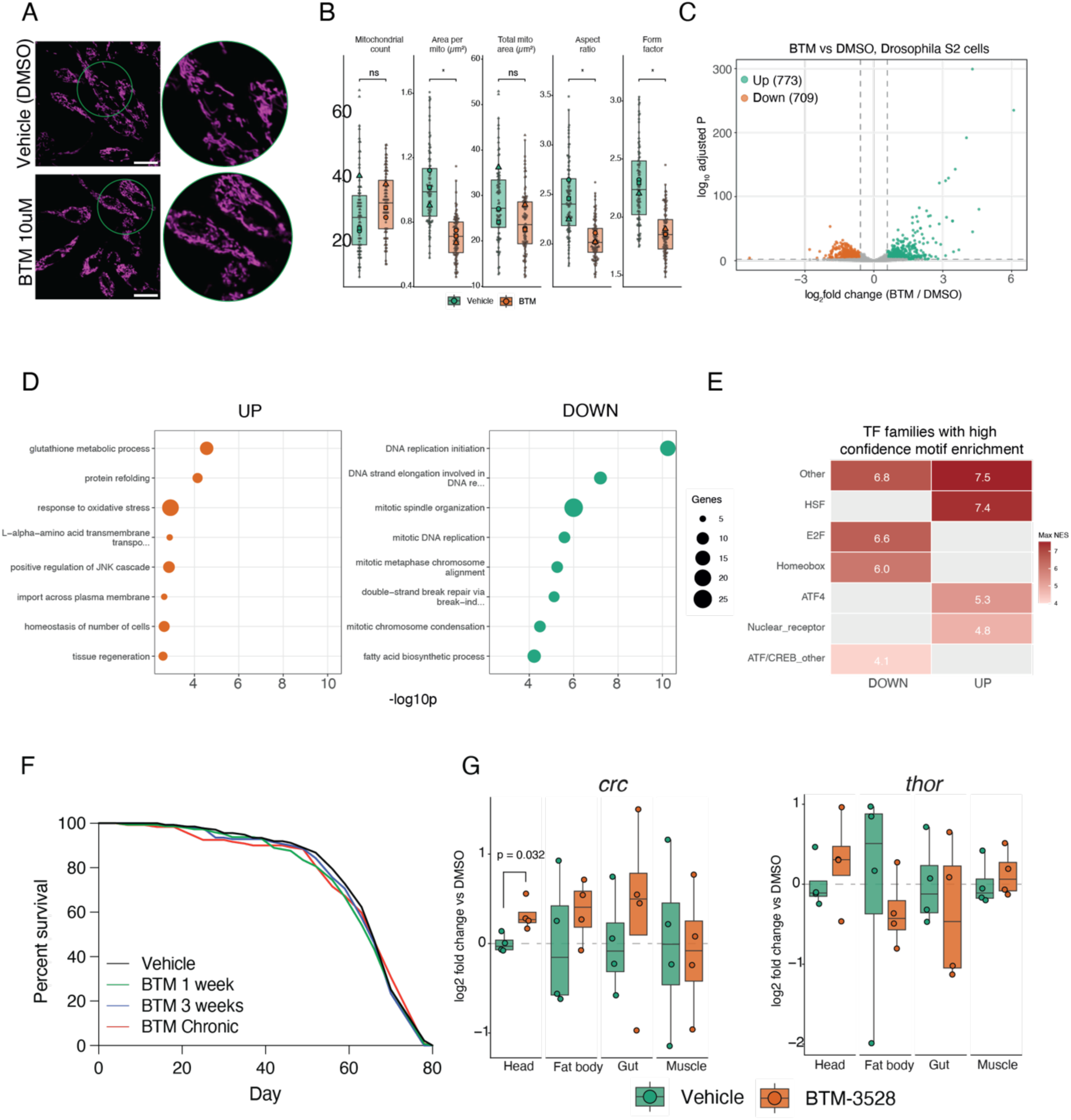
**a)** Representative images of S2R+ cells exposed to either vehicle or 10μM BTM compound for 4h, marked with MitoTracker Green. **b)** Superplots with the quantifications of Mitochondrial count, Area, Mitochondrial area, Aspect Ratio and Form Factor and of cells treated with 10μM BTM compound for 4h. **c)** Volcano plot of differentially expressed genes in S2R+ cells treated with 10 µM BTM-3528 for 6 h compared with vehicle-treated controls (n = 5 vehicle, 3 BTM). Dashed lines show the significance thresholds (FDR < 0.01 and |log2FC| > 0.585). **d)** GO biological process terms enriched among genes up-(top) and downregulated (bottom) by BTM. Top 8 terms per direction among those at classic Fisher p < 0.05, ranked by the weight01 algorithm. **e)** Transcription factor binding site (TFBS) motif enrichment among genes up- and downregulated by BTM, summarised as the maximum normalised enrichment score (NES) per TF family. Families with at least one motif at NES ≥ 4 are shown; blank tiles indicate no motif above this display threshold. Enrichment was called at NES ≥ 3. **f)** Lifespan of *wDah* wild-type females upon exposure to vehicle (DMSO) or BTM compound (10uM) at different periods of early adulthood. Vehicle n = 132 dead/7 censored, BTM 1 week n = 141 dead/6 censored, BTM 3 weeks n = 138 dead/2 censored, BTM chronic during adulthood n = 117 dead/6 censored, no significant differences were observed in any comparisons, log-rank test. **g)** *crc* and *thor* expression in dissected organs after 24 h of feeding 10 µM BTM, relative to DMSO controls and normalised to the mean of *tubulin* and *actin*. Box plots show median and quartiles; dots show individual samples (n = 4 per condition per tissue). P-values are from a generalised least-squares model with tissue-specific variances and RNA batch as a covariate (BTM versus DMSO within each tissue, Satterthwaite degrees of freedom); p-values < 0.05 are shown. Fold changes, 95% CIs and p-values for all organs are given in Table S5.

Encouraged by the similarity between the transcriptional responses induced by BTM in cells and ATF4 in flies *in vivo*, we fed adult flies with BTM compound. BTM treatment length was the same as the ATF4 inductions and recoveries performed in vivo. However, BTM treatment did not change lifespan (Figure 5F), nor did it reduce body weight or egg laying; instead BTM-fed flies were slightly heavier at several timepoints (Figure S6B and C). To test target engagement in the four major tissues, we measured by qPCR, in individual organs after 24 h of exposure, *crc* and *thor*, two of the genes most strongly induced by ATF4 and by BTM in cells (Table S5). *Crc* was modestly increased in the head (1.24-fold, p = 0.03), but not in the fat body, gut or muscle, and *thor* was unchanged in all four organs (Figure 5G). Altogether, these data show that BTM elicits mitochondrial fragmentation and an ATF4-associated transcriptional response in *Drosophila* cells, but that feeding achieved only limited target engagement in vivo, insufficient to extend lifespan.

### Transient neuronal ATF4 activation drives non-autonomous chromatin remodelling and transposon gene expression changes in the aged fat body

Neuronal ATF4^on-off^ therefore remains the most effective way to harness the benefits of ATF4, and we used it to ask what a single week of neuronal ATF4 activity leaves behind. To identify its long-term consequences in peripheral organs, we profiled the transcriptomes of head, gut, thoracic muscle and fat body in 50-day-old neuronal ATF4^on-off^ flies (41 days after switching ATF4 off), an age at which they are old, but mortality has not yet risen sharply. We also profiled uninduced, driver-alone controls to separate ATF4-driven changes from responses to RU_486_ itself. RU_486_ alone produced a substantial transcriptional response, most pronounced in gut and head, but this response was essentially independent of the ATF4 response genome-wide (Table S3): only 29 genes across all four tissues were significant in both genotypes, 19 of them changing in opposite directions. Removing genes that changed in the same direction in controls retained 93 to 100% of ATF4 differentially expressed genes in every tissue (Table S3).

Of the four major tissues analysed, the fat body showed by far the strongest response, with 3014 differentially expressed genes compared with fewer than 110 in head, gut or muscle (Figure 6A). GO analysis in the fat body revealed two opposing signatures: coordinated upregulation of chromatin-related processes, namely chromatin organisation, heterochromatin formation and histone deacetylase activity, alongside downregulation of metabolic processes, most prominently mitochondrial oxidative phosphorylation genes (Figure S7A). KEGG gene set enrichment analysis reproduced the same pattern, with enrichment of ATP-dependent chromatin remodelling and the Polycomb repressive complex, while showing decreased oxidative phosphorylation (Figure S7B). The induction was broad across the heterochromatin pathway: of 99 genes annotated to heterochromatin formation, 39 were significantly upregulated, including histone methylation regulators (G9a, Hmt4-20, Su(var)3-3), HP1b, Polycomb group genes (Pc, Psc, esc) and deacetylases (HDAC1, Sirt1, Sirt6) (Figure 6B).

**Figure 6.**
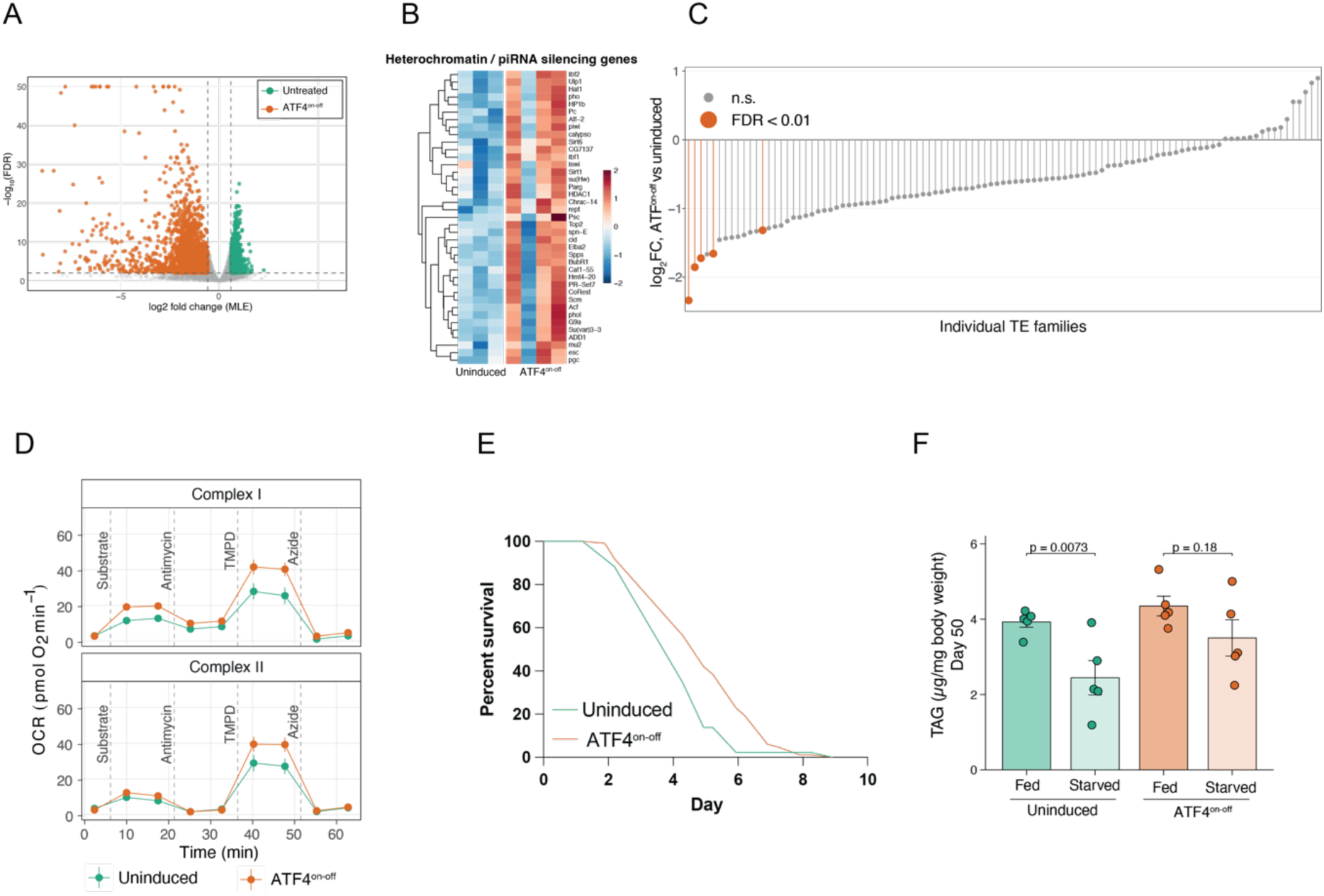
**a)** Volcano plot of differentially expressed genes in the fat body of 50-day-old neuronal ATF4^on-off^ flies versus uninduced controls. Dashed lines show the significance thresholds (FDR < 0.01 and |log2FC| > 0.585). **b)** Heterochromatin and piRNA pathway genes significantly upregulated in the fat body of neuronal ATF4^on-off^ flies (FDR < 0.01, |log2FC| > 0.585). Heatmap of row-scaled (z-score) variance-stabilised expression in each biological replicate. **c)** Change in expression of individual transposable element (TE) families in the fat body of 50-day-old neuronal ATF4^on-off^ flies, ordered by log_2_FC; highlighted families reach FDR < 0.01. 87/103 families decreased; total TE burden was 0.51-fold of controls (Welch’s t-test p = 0.064, n = 3 vs 4 pools). Values for all families are given in Table S3. **d)** Oxygen consumption traces of frozen fat body homogenates from 50-day-old neuronal ATF4^on-off^ flies and uninduced controls. Complex I was fed NADH and complex II succinate plus rotenone, followed by sequential injections of antimycin A, TMPD/ascorbate and azide (dashed lines). Mean ± SEM of n = 4 cohorts, technical wells averaged within each cohort; wells showing substrate depletion are shown but were excluded from complex II quantification. e) Starvation survival (1% agarose) of 50-day-old neuronal ATF4^on-off^ flies. Uninduced n = 71 dead/20 censored, ATF4^on-off^ n = 96 dead/20 censored; p = 0.002, log-rank test. f) Whole-body triacylglycerol (TAG) normalised to body weight in 50-day-old neuronal ATF4^on-off^ flies and uninduced controls, fed or after 72 h of starvation (1% agarose). Bars show mean ± SEM of n = 5 samples of 3 flies per condition; dots show individual samples. Statistics are from a linear model on log-transformed values: brackets show fed versus starved within each condition (*emmeans* contrasts); the RU486 × starvation interaction tests whether the proportional loss differs between conditions (ratio of fraction retained, ATF4^on-off^/uninduced = 1.35, 95% CI 0.79–2.29, p = 0.25).

Transposable element (TE) de-repression is a hallmark of ageing. Heterochromatin silencing of TEs becomes less effective with age, and reducing TE abundance can extend lifespan^46,47^. Given the coordinated increase of heterochromatin genes, we asked whether TE expression was also reduced. Quantifying our fat body libraries against a combined transcript and transposon index, normalised using gene-level size factors so that a global shift could not be removed by normalisation (see Methods), we found that total TE burden tended to be lower in the fat body of neuronal ATF4^on-off^ flies, with 87 of 103 TE families shifting downward (Figure 6C), and RT-qPCR confirmed reduced expression of *gypsy*, one of the most abundant TE families (Figure S7C). In contrast, TE expression did not shift in driver-alone controls (Figure S7D).

These results show that, despite that the fat body never expressed the ATF4 transgene, it carried the strongest and most durable transcriptional signature of the intervention. Transient ATF4 activation in neurons therefore acts non-autonomously to establish a lasting chromatin and metabolic state in a distant tissue, weeks after the stimulus has ended.

### Neuronal ATF4^on-off^ increases OXPHOS capacity in the aged fat body

The reduction in OXPHOS gene expression in old fat bodies prompted us to ask whether neuronal ATF4 activation leaves a lasting functional imprint on fat body mitochondria. We measured respiratory chain capacity in frozen fat body homogenates from 50-day-old females using a XFe96 Analyzer. Contrary to the transcriptional data, respiratory capacity was higher in neuronal ATF4^on-off^ flies. Complex I capacity was almost two-fold elevated, with a similar trend for complex II (Figure 6D, S8A). The same pattern was present after ubiquitous induction followed by recovery, although the differences did not reach significance (*DaGS*, Figure S8A and B). Higher respiratory capacity could reflect either more mitochondria or more active ones. To distinguish these possibilities, we measured levels of ETC subunits by western blot, normalised to either tubulin or vinculin. Neither COX4 (complex IV) nor ATP5A (complex V) was increased in old fat bodies (Figure S8C). This suggests that the observed enhancement in complex I (and complex II) respiratory capacity reflects intrinsic mitochondrial functional optimization or higher ETC efficiency, rather than a mere expansion of total mitochondrial mass.

Consistent with a lasting functional remodelling of the fat body, the main energy-storage organ of the fly, 50-day-old neuronal ATF4^on-off^ flies were more resistant to starvation than age-matched controls (Figure 6E, S8D). To test whether this reflected altered use of energy reserves, we measured whole-body TAG in fed flies and after 72h of starvation. Starvation significantly reduced TAG in control flies (58% retained, p = 0.0073), whereas the reduction in neuronal ATF4^on-off^ flies did not reach significance (78% retained, p = 0.18, Figure 6F). The difference between conditions was not significant (p = 0.25, LM), but body weight showed the same pattern (80% versus 92% retained, p = 0.058; Figure S8E). These data suggest that neuronal ATF4^on-off^ flies draw less on their energy reserves during starvation.

Together, these results show that a single week of neuronal ATF4 activation leaves the aged fat body in a distinct functional state, with greater respiratory capacity, sparing their energy reserves. The benefit of this intervention therefore appears to reside in a remodelled, non-autonomous metabolic state rather than in the acute stress response itself.

## Discussion

Stress responses are widely assumed to be pro-longevity^48–50^, but the ISR has resisted that generalisation, extending lifespan in some settings and shortening it in others. Our findings help reconcile conflicting reports on the role of ATF4 in ageing and disease, spanning from yeast to mammals^10,11,13,38,51^. The apparent contradiction extends to single tissues: in the fly brain, ATF4 has been reported both to cause neurodegeneration^18^ and to prevent it^19,20^. Rather than reflecting species or model differences, these outcomes are consistent with a single pathway whose effect is set by the intensity, duration and location of its activation.

Our results also bear on the recent finding that suppression, rather than activation, of ATF4 extends lifespan^38^. We confirmed that chronic ATF4 activation throughout adulthood shortened lifespan. Transient activation confined to early adulthood, by contrast, had the opposite effect, indicating that the sign of the outcome depends on how long the pathway is engaged rather than on whether it is engaged at all. The duration of activation was decisive, but the optimum differed between tissues: three weeks was more effective than one when ATF4 was activated ubiquitously, whereas in neurons, only one week expression extended lifespan. The window of sensitivity was not restricted to early adulthood, as midlife activation also extended lifespan, although less markedly. This fits a hormetic interpretation, in which a brief stimulus elicits an adaptive response while a prolonged one becomes a burden, with that threshold reached sooner in neurons than across the whole animal, and probably promoting longevity through different mechanisms.

The fact that the fat body, rather than any tissue closer to the stimulus, carries the strongest signature of a neuronal intervention is the clearest indication that neuronal ATF4 acts non-autonomously. Two features of this signature are notable. First, the signature is directional with respect to ageing: heterochromatin loss and transposable element de-repression are conserved features of old age^46,52^, and the fat body of old ATF4^on-off^ flies showed the opposite pattern, with induction of heterochromatin genes and lower TE expression. Second, the effects of a one-week pulse in early adulthood are still detected in a different tissue six weeks later, which argues for a self-sustaining mechanism rather than a slowly decaying signal. Whether changes in chromatin structure cause the persistence or merely accompany it remains to be tested. The mitochondrial arm of this signature behaved differently. Respiratory chain capacity in old fat bodies was higher without an increase in ETC subunit abundance, indicating a change in the activity rather than the amount of the respiratory machinery. Transcript abundance of the OXPHOS machinery was reduced in the same tissue, but respiratory flux is set principally by substrate supply, supercomplex assembly and membrane potential rather than by subunit mRNA levels, so the two measurements are not expected to track one another and we do not interpret their divergence as requiring a single explanation. One possibility is that these flies retain better-functioning mitochondria into old age and therefore sustain capacity with less transcriptional investment, consistent with the long half-lives reported for mitochondrial proteins^53^. Investigating this will require measurements of mitochondrial content and turnover.

Another important question is the identity of a brain-derived signal. Inter-organ communication from the nervous system is well established in *Drosophila* ^54–56^, and FGF signalling was an attractive candidate given its links to stress signalling and metabolic regulation^57,58^, but our data do not support it: neither *branchless* nor its receptors were altered at either timepoint. In our ubiquitous paradigm, heads during induction showed increased *Ilp3* and *Ilp5*, insulin-like peptides (Table S2) produced by brain insulin-producing cells, as well as *Ilp6* and *CCHa2*, which are established regulators of systemic metabolism and lifespan.^59–61^. However, the increase we see is in the opposite direction to that expected for a pro-longevity signal, although transcript levels do not report peptide release, which is controlled post-transcriptionally. Whether any of these ligands mediates the fat body response remains to be tested.

Our motif analysis suggests that ATF4 engages different partners depending on the tissue and the phase of the response^62,63^. During induction, ATF4 motifs were accompanied by HSF motifs in head, gut and muscle, and by C/EBP motifs in muscle and gut, consistent with ATF4 heterodimerisation with C/EBP family members^64,65^. HSF1 is a plausible contributor to the benefits we observe, having been linked to neuronal survival and stress resistance^66^, and in *C. elegans*, lifespan extension conferred by ATF4 requires HSF-1^10^. The overlap between tissues supports this two-phase view. During induction, all tissues shared a common core of ATF4 targets, whereas after recovery the response became largely tissue-specific. The persistence of oxidative stress resistance long after ATF4 and its acute signature have resolved therefore cannot be explained by continued ATF4 activity. We favour a relay model, in which the acute ATF4 response initiates a transcriptional state that is subsequently maintained by other factors, such as SOX or GATA factors. A comparable paradigm may operate for other transcription factors: we recently found that transient dFOXO activation in the same window of early adulthood also promotes longevity^5^, though from a different organ and through different pathways, suggesting that several independent routes converge on durable late-life benefits.

Furthermore, the response of *Drosophila* cells to BTM points to deep conservation of this pathway despite flies lacking several of its molecular players. Flies lack clear orthologues of several components that mediate the mitochondrial stress response in mammals: there is no evident OMA1 orthologue, although Rhomboid*-7* has been proposed to perform its role in controlling mitochondrial morphology^67,68^. Also, flies lack HRI, which together with DELE1 relays mitochondrial stress to eIF2α in mammals^69,70^. The fact that BTM nonetheless elicits mitochondrial fragmentation and a transcriptional response enriched for ATF4 and HSF targets in fly cells suggests that an equivalent relay exists, perhaps through PERK or GCN2, whose downstream response is also mediated by ATF4^18–20^. The weaker response *in vivo* has several possible explanations: the compound may be degraded or poorly absorbed in food, 24h may be too short for a response to develop in adult tissues, or the dose reaching peripheral organs may be insufficient to have beneficial effects. It is notable that the only tissue showing a response was the head, rather than the gut or fat body, which are in more direct contact with the food.

In conclusion, our study establishes ATF4 as a programmable determinant of longevity whose outcome depends on when, for how long, and in which tissue it is engaged. A single week of neuronal ATF4 activity in early adulthood is sufficient to establish lasting systemic adaptations, without the metabolic and reproductive costs of whole-body activation. The benefit of a stress response lies not only in its duration, but also in where and when it occurs.

### Limitations of the study

Our study establishes that neuronal ATF4 activation is sufficient to extend lifespan and healthspan, but does not address whether it is required for the effects of ubiquitous activation; the different phenotypic profiles of the two paradigms suggest that they act, at least in part, through distinct mechanisms. The brain-derived signal that mediates the fat body response is still to be identified, and our neuronal transcriptomic dataset was collected only at old age, so changes occurring during the induction window remain uncharacterised. The chromatin and transposable element changes in the fat body are correlative, and whether they cause the persistence of the phenotype remains to be tested. Finally, as for many longevity interventions in *Drosophila*, our lifespan effects were restricted to females, and the basis of this sex specificity is unknown.

## Author contributions

Conceptualization: G.M.C.; Methodology: P.S.D., M.L., G.M.C.; Investigation: P.S.D., Ó.T.F., S.S.R., O.M.; Formal analysis: Ó.T.F., P.S.D., G.M.C.; Visualization: P.S.D., G.M.C.; Resources: M.L., G.M.C; Supervision: G.M.C.; Funding acquisition: G.M.C.; Writing: original draft: G.M.C.; Review & editing: all authors.

## Competing interests

M.L. is a co-founder of Enspire Bio LLC, and a consultant of Capacity-Bio, and received sponsored Research agreement from Bantam Pharmaceutical in the past.

## Acknowledgments

We thank Hyung Don Ryoo, Joseph Bateman, Nazif Alic, Linda Partridge, Marco Milán for providing fly stocks, and Marta Llimargas and Jordi Casanova for also providing fly stocks and fruitful discussions improving the manuscript. We also thank Jordi Bernués for providing cell stocks and technical assistance. We also thank Dr. Enric Ureña for critical comments on the manuscript. Finally, we thank the Mitochondria and Metabolism Platform (MMP-IBMB) for technical assistance. We thank Matthew Kostura for the kind gift of BTM-3528. This study was supported by a Ramón y Cajal Fellowship (RYC2021-031912-I) and PID2023-146293NA-I00 from MCIN/AEI of the “ERDF A way of making Europe”, by the European Union, both awarded to G.M.C. M.L. is funded by PID2021-127278NB-I00 from MCIN/AEI.

## Supplementary figures

**Figure S1.**
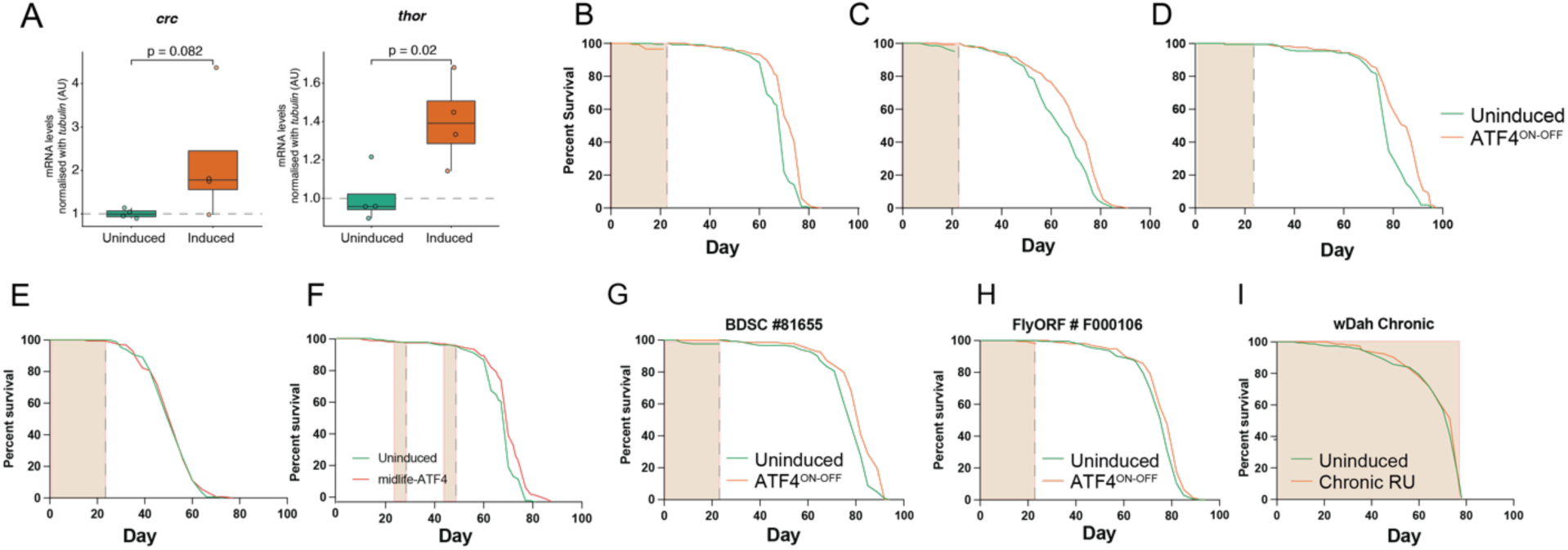
**a)** qPCR of whole-body extracts of heads of female DaGS>UAS-ATF4 flies, 1 week induced. For *crc* p = 0.082, for *thor* p = 0.02. **b-d)** Additional independent replicates of ubiquitous ATF4^on-off^ experiments. b: Uninduced n = 122 dead/20 censored, ATF4^on-off^ n = 137 dead/11 censored, p = 6.02 x 10^-7^, log-rank test. c: Uninduced n = 136 dead/15 censored, ATF4^on-off^ n = 134 dead/5 censored, p = 1 x 10^-4^, log-rank test. d: Uninduced n = 160 dead/20 censored ATF4^on-off^ n = 151 dead/18 censored, p = 2 x 10^-9^, log-rank test. **e)** Survival of male ATF4^on-off^ flies. Uninduced n = 151 dead/19 censored, ATF4^on-off^ n = 136 dead/32 censored, p = 0.57, log-rank test. **f)** Midlife ATF4^on-off^ lifespan. Uninduced n = 122 dead/20 censored, midlife ATF4^on-off^: 150 dead/7 censored, p = 0.0003, log-rank test. **g)** ATF4^on-off^ experiment employing a second, independent UAS-ATF4 line (BDSC #81655). Uninduced n = 152 dead/17 censored, ATF4^on-off^: 136 dead/16 censored, p = 5.42 x 10^-6^, log-rank test. **h)** ATF4^on-off^ experiment employing a third, independent UAS-ATF4 line (FlyORF #F000106). Uninduced n = 173 dead/7 censored, ATF4^on-off^: 156 dead/9 censored, p = 0.01, log-rank test. **i)** Chronic RU exposure in *DaGS* > *wDah* driver-alone experiment. Uninduced n = 154 dead/3 censored, ATF4^on-off^: 146 dead/4 censored, p = 0.49, log-rank test Shaded areas in lifespan curves indicate the times of induction of ATF4 by addition of RU^486^ to the fly food.

**Figure S2.**
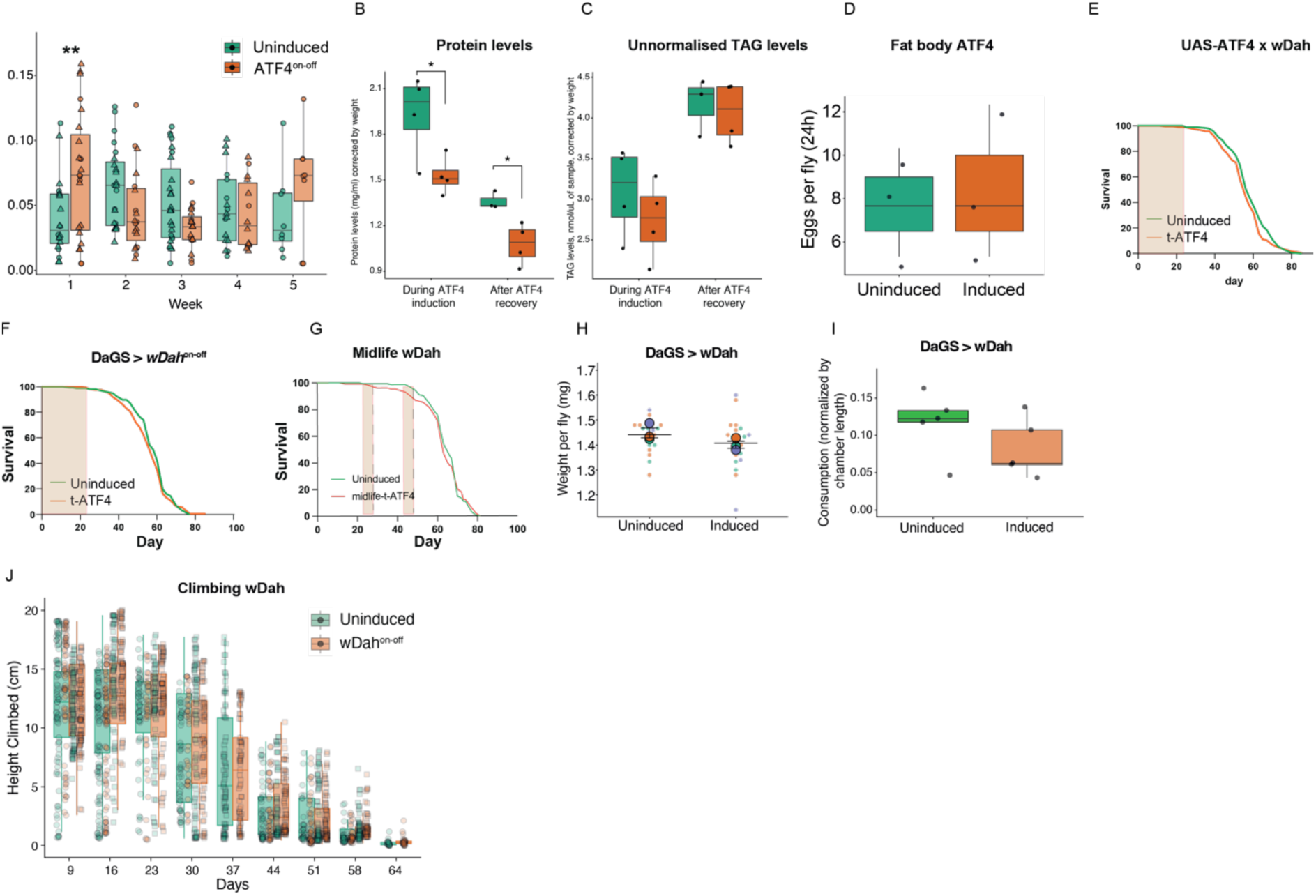
**a)** Lifetime volume of food consumed by ATF4^on-off^ flies over a 5-week period, measured by CAFE assay (n = 10 individual flies per condition, 2 biological replicates). Consumption was normalized by evaporation controls. Data shown as mean ± SEM. ** p < 0.01 at week 1, no significant effect of RU^486^ induction detected at any other timepoint, linear mixed effects model with replicate as a random effect, emmeans pairwise comparisons with Tukey adjustment. **b)** BCA quantifications of ATF4^on-off^ flies at day 9 (during ATF4 induction) and day 35 (after ATF4 recovery), normalised by total body weight. * Indicates p < 0.05, student t-test. **c)** TAG levels of ATF4^on-off^ flies at day 9 (during ATF4 induction) and day 35 (after ATF4 recovery), normalised by total body weight. No significant differences detected, student t-test. **d)** Egg counts of *Lsp2*-GS > UAS-ATF4, one week old, one week induced, over a 24h period (n = 3 vials per condition). **e)** ATF4 construct-alone control of ATF4^on-off^ experiments, without the DaGS construct. Uninduced n = 172 dead/5 censored, ^on-off^ 3W n = 164 dead/9 censored, p = 0.091, log-rank test. **f)** Driver alone control of ATF4^on-off^ experiments, without the ATF4 construct. Uninduced n = 156 dead/7 censored, DaGS^on-off^ n = 158 dead/10 censored, p = 0.29, log-rank test. **g)** Driver-alone control of midlife-ATF4 experiments, without the ATF4 construct. Uninduced n = 137 dead/10 censored, midlife-switch n = 129 dead/6 censored, p = 0.84, log-rank test. **h)** Body weight of 9-day-old DaGS > *wDah* driver-alone flies after 1 week of RU_486_, compared with uninduced controls. Small dots show individual samples coloured by replicate; large dots show replicate means; bar shows mean ± SEM of replicate means (n = 4-10 samples per condition per replicate, 3 biological replicates). RU p = 0.21, linear mixed-effects model with replicate as a random effect. **i)** Volume of food consumed by driver alone controls of DaGS > *wDah* experiments measured by CAFE assay (n = 5 individual flies per condition). Consumption was normalized by evaporation controls. **j)** Related to Figure 2E, height climbed by *wDah*^on-off^ flies vs uninduced controls, from 2 independent trials. n = 88-108 per experimental condition. Boxplots: quartiles; whiskers: extremes; overlay: individual data points. Effect of t-RU p = 0.6719, age p < 0.0001, age-by-t-RU interaction p = 0.2873, mixed-effects LM. In e, f and g, shaded orange area represents the period of RU exposure.

**Figure S3.**
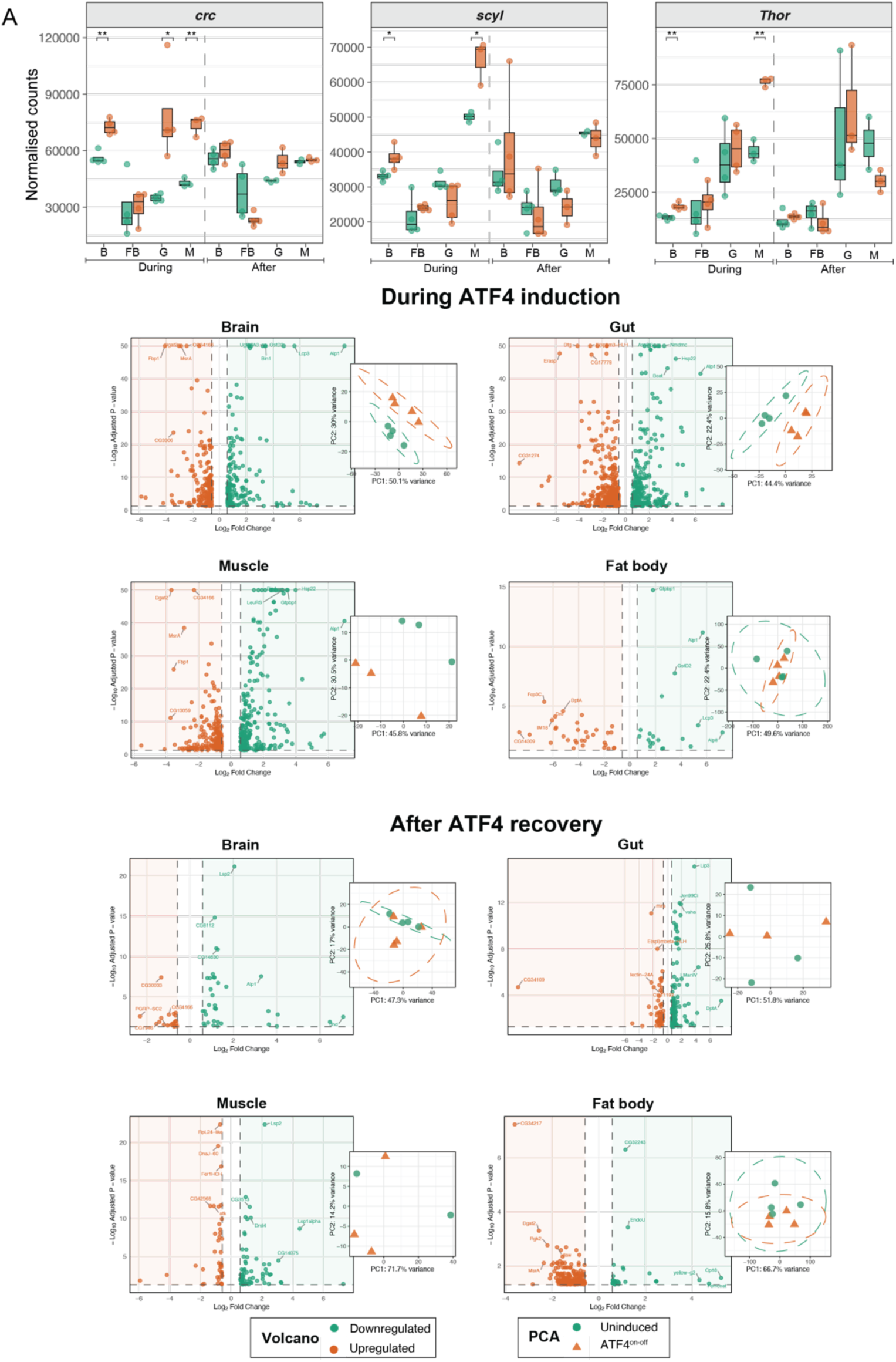
**a)** Normalised counts of RNAseq samples for *crc* and its targets *scyl* and *thor,* during ATF4 induction (day 9, after 7 days of RU486) and after recovery (day 35, 12 days after RU486 withdrawal) * indicates p-value < 0.05, ** = p-value < 0.01 and *** = p-value < 0.001, student t-test. **b)** Volcano plots of differentially expressed genes from the conditions shown in Figure 3A. Principal component analysis (PCA) from the VST-transformed gene expression data, from the conditions shown in Figure 3A. PC1 and PC2 are shown, with their proportion of total variance shown in the axis.

**Figure S4.**
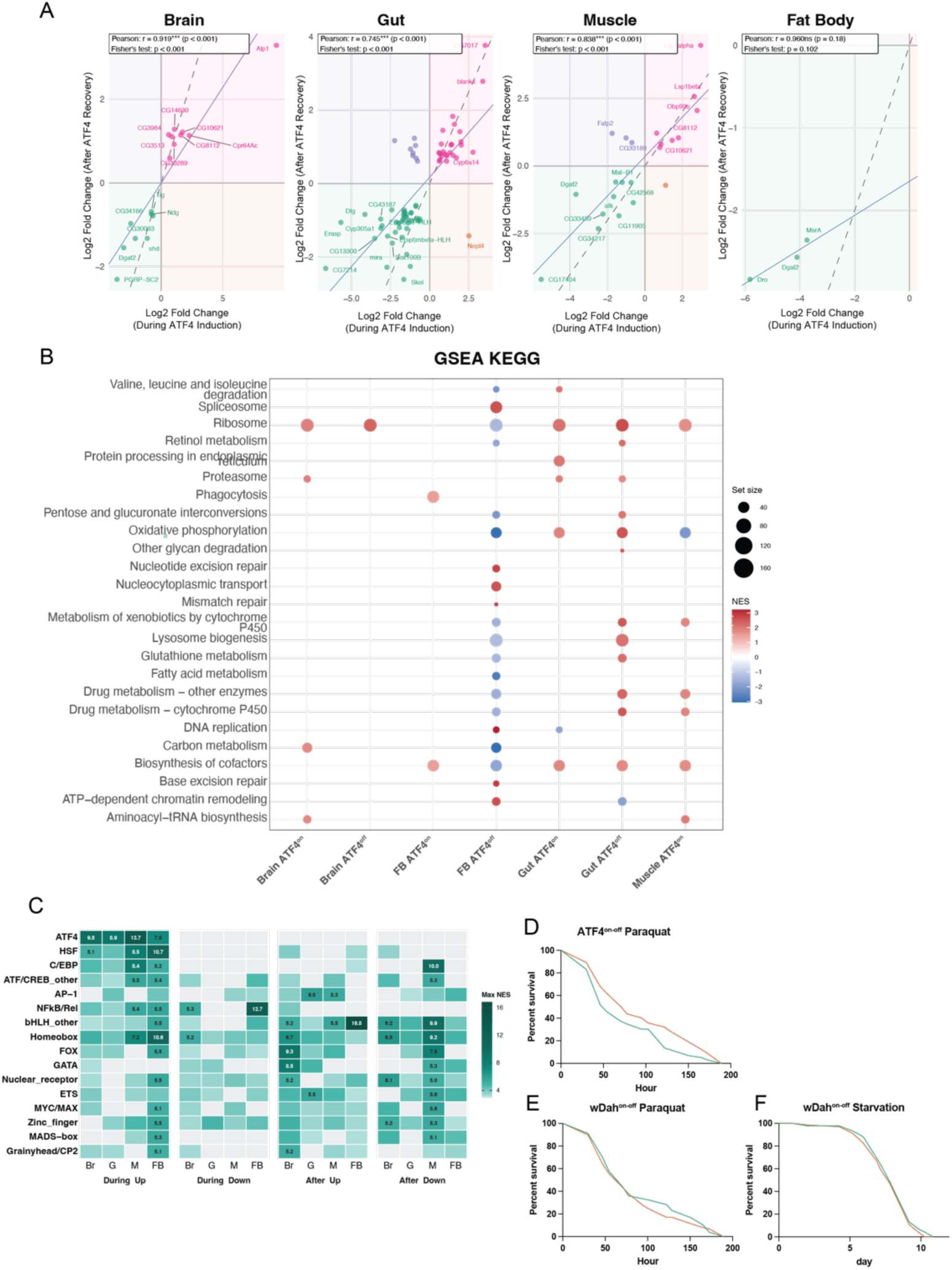
**a)** Correlation plots of the overlapping DE genes across tissues between timepoints. Pearson correlations and overlap statistics (Fisher’s exact test) are shown in each graph. The direction of shared genes was tested with a two-sided binomial test against 50% concordance. Head: 19 shared vs 1.8 expected (OR = 15.3, padj = 1.2 × 10⁻¹⁴; 19/19 same direction, padj = 5.7 × 10⁻⁶). Gut: 88 shared vs 13.7 expected (OR = 11.4, padj = 6.4 × 10⁻⁴⁹; 78/88 same direction, padj = 1.0 × 10⁻¹³). Muscle: 13 shared vs 1.2 expected (OR = 16.4, padj = 1.1 × 10⁻¹⁰; 12/13 same direction, padj = 3.4 × 10⁻³). Fat body: no shared genes (0.1 expected; padj = 1). **b)** KEGG pathway analysis of the different strata of ATF4^on-off^, showing the Normalised Enrichment Score (NES), and the size of each KEGG pathway. **c)** Heatmap showing the maximum Normalised Enrichment Score (NES) for transcription factor binding site (TFBS) motifs within each TF family. Each column represents one tissue (Br, brain; G, gut; M, muscle; FB, fat body) within each condition group (During Up/Down: During ATF4 induction timepoint upregulated/downregulated ATF4-specific genes; After Up/Down: After ATF4 recovery). Only TF families reaching NES ≥ 5 in at least 2 conditions are shown. Tile colour reflects the highest NES across all motifs annotated to that family; grey tiles indicate no enrichment above the analysis threshold (NES = 3). Numbers indicate Max NES values ≥ 5. **d)** Second replicate of the assay against oxidative stress (20mM paraquat), uninduced n = 132 dead/2 censored, ATF4^on-off^ n = 112 dead/4 censored. P = 0.004, log-rank test. **e)** Driver-alone control of the assay against oxidative stress (20mM paraquat), with no ATF4 transgene. uninduced n = 78 dead/2 censored, *wDah*^on-off^ n = 88 dead/2 censored. P = 0.63, log-rank test. **d)** Starvation survival assay of *wDah* driver-alone controls (1% agarose). Uninduced n = 136 dead/0 censored, *wDah*^on-off^ n = 138 dead/1 censored. P = 0.22, log-rank test.

**Figure S5.**
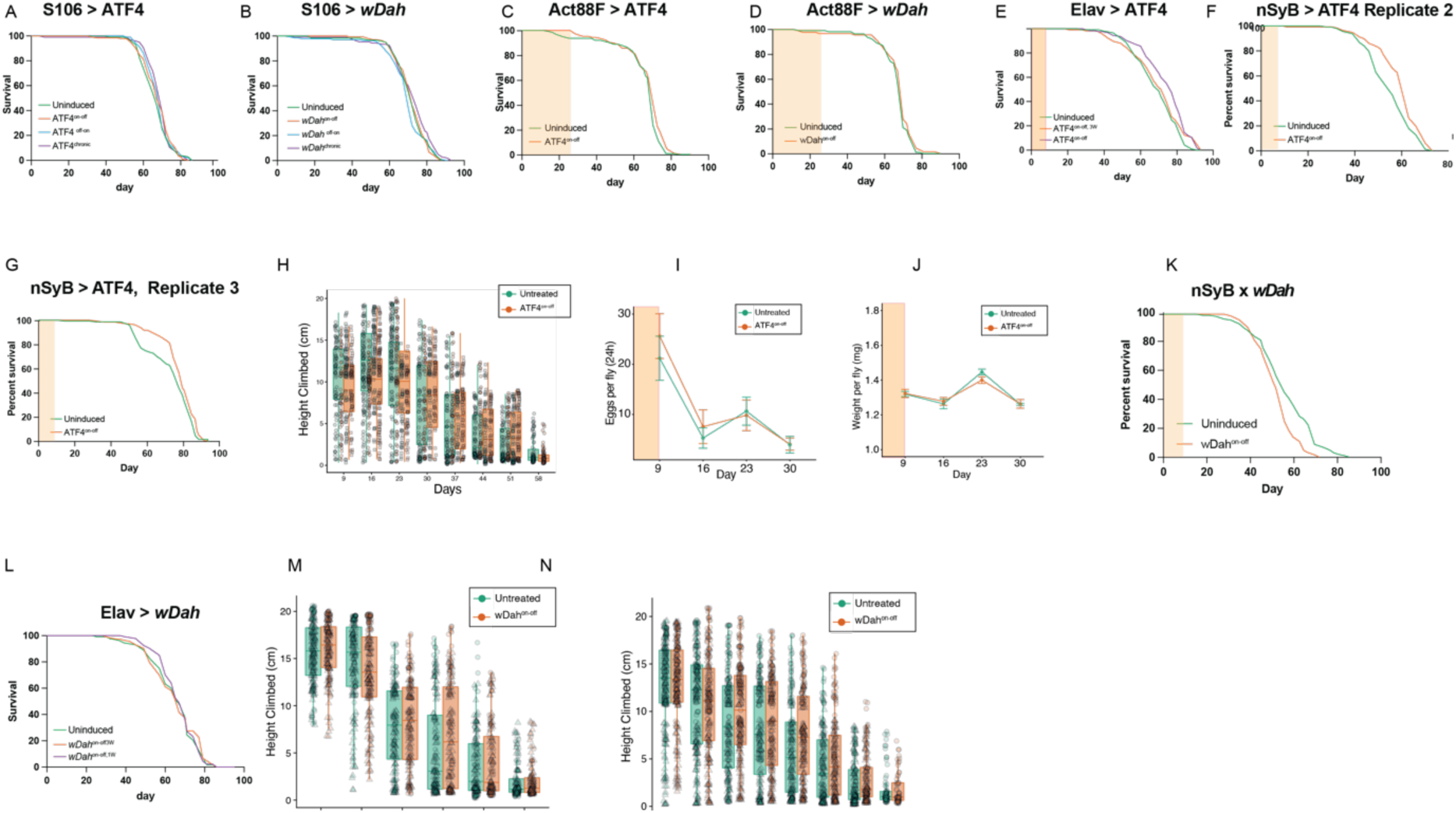
**a)** Lifespan curves of S106 > UAS-ATF4, at the different times shown. None of the comparisons between conditions reached statistical significance (p>0.05 log-rank test). Uninduced n = 129 deaths / 4 censored, ATF4^on-off^ (from day 2 to day 23) n = 111 dead/7 censored, ATF4^off-on^ (day 23 until death) n = 100 dead / 2 censored and ATF4^chronic^ n = 96 dead / 8 censored. **b)** S106, driver-alone control experiment. Uninduced n = 118 deaths / 13 censored, *wDah*^on-off^ (from day 2 to day 23) n = 134 dead/11 censored, *wDah*^off-on^ (day 23 until death) n = 133 dead / 13 censored and *wDah*^chronic^ n = 103 dead / 19 censored. **c)** Muscle (Act88F-GS) ATF4^on-off^ lifespan. Uninduced n = 125 deaths / 5 censored, ATF4^on-off^ (from day 2 to day 23) n = 106 dead/4 censored. P > 0.05, log-rank test. **d)** Muscle driver-alone control Uninduced n = 109 deaths / 6 censored, *wDah*^on-off^ (from day 2 to day 23) n = 89 dead/6 censored. **e)** Additional neuronal (*Elav-*GS) driver lifespan. Uninduced n = 117 dead/2 censored, ATF4^on-off^ 1W (day 2 until day 9) n = 125 dead/4 censored, ATF4^on-off^ 3W (day 2 until day 23) n = 116 dead/4 censored. Uninduced vs ATF4^on-off^1W p = 8.73 × 10⁻⁵; uninduced vs ATF4^on-off^ 3W p = 0.083, log-rank test. **f)** Second replicate of neuronal ATF4^on-off^ with the *nSyb-*GS driver. Uninduced n = 150 dead / 7 censored, ATF4^on-off^ ^1W^ (day 2 until day 9) n = 133 dead / 13 censored. Uninduced vs ATF4^on-off^ ^1W^ p = 2.5 x 10^-3^ log-rank test. **g)** Third replicate of neuronal ATF4^on-off^ with the *nSyb-*GS driver. Uninduced n = 92 dead / 22 censored, ATF4^on-off^ ^1W^ (day 2 until day 9) n = 103 dead / 36 censored. Uninduced vs ATF4^on-off^ ^1W^, p = 2.4 x 10^-4^ log-rank test. **h)** Height climbed by neuronal ATF4^on-off^ (*Elav*-GS) flies vs uninduced controls, pooled from 2 independent trials. Boxplots: quartiles; whiskers: extremes; overlay: individual data points, shaped by trial. Effect of RU486 p < 0.0001, age p < 0.0001, RU:age interaction p < 0.0001; RU:age:genotype interaction versus driver-alone controls (panel n) p = 0.0135; linear mixed-effects model with flipper as a random effect and trial as a covariate. n = 2460 observations after trimming. **i)** Lifetime fecundity of neuronal ATF4^on-off^ (Elav-GS) and uninduced flies, eggs per female per 24 h. n = 5 vials of 3-5 females per condition per day. No significant differences (Welch’s t-test per day, all p > 0.50). **j)** Lifetime weight measurements of whole flies, comparing neuronal ATF4^on-off^ (*Elav-*GS) vs uninduced controls. n = 6-22 vials of 2-13 flies per condition; day 9 pooled from 3 replicates. No significant differences (Welch’s t-test per day, all p > 0.08) **k)** Driver-alone control for *nSyb-*GS lifespan. Uninduced n = 122 dead / 29 censored, *wDah*^on-off^ (day 2 until day 9) n = 125 dead / 57 censored. p = 1.3 x 10^-3^ log-rank test. **l**) *Elav*-GS driver-alone control lifespan. Uninduced n = 126 dead/2 censored, *wDah*^on-off^ ^1W^ (day 2 until day 9) n = 133 dead/1 censored, wDah^on-off^ ^3W^ (day 2 until day 23) n = 110 dead/7 censored. No significant comparisons (p = 0.75 and p = 0.56), log-rank test. **m)** Related to Figure 4B. Height climbed by neuronal *wDah*^on-off^ flies (*nSyb* driver) vs uninduced controls, from 2 independent trials. n = 122-183 per experimental condition. Boxplots: quartiles; whiskers: extremes; overlay: individual data points. Effect of RU p = 0.92, age p < 0.0001, age-by-RU interaction p = 0.2526, mixed-effects LM. **n)** Height climbed by neuronal *wDah*^on-off^ flies (*Elav* driver) vs uninduced controls, from 2 independent trials. n = 61-208 per experimental condition. Boxplots: quartiles; whiskers: extremes; overlay: individual data points. Effect of RU p = 0.508, age p < 0.0001, age-by-RU interaction p = 0.1147, mixed-effects LM. All the source data is shown in Supplementary Tables X.

**Figure S6.**
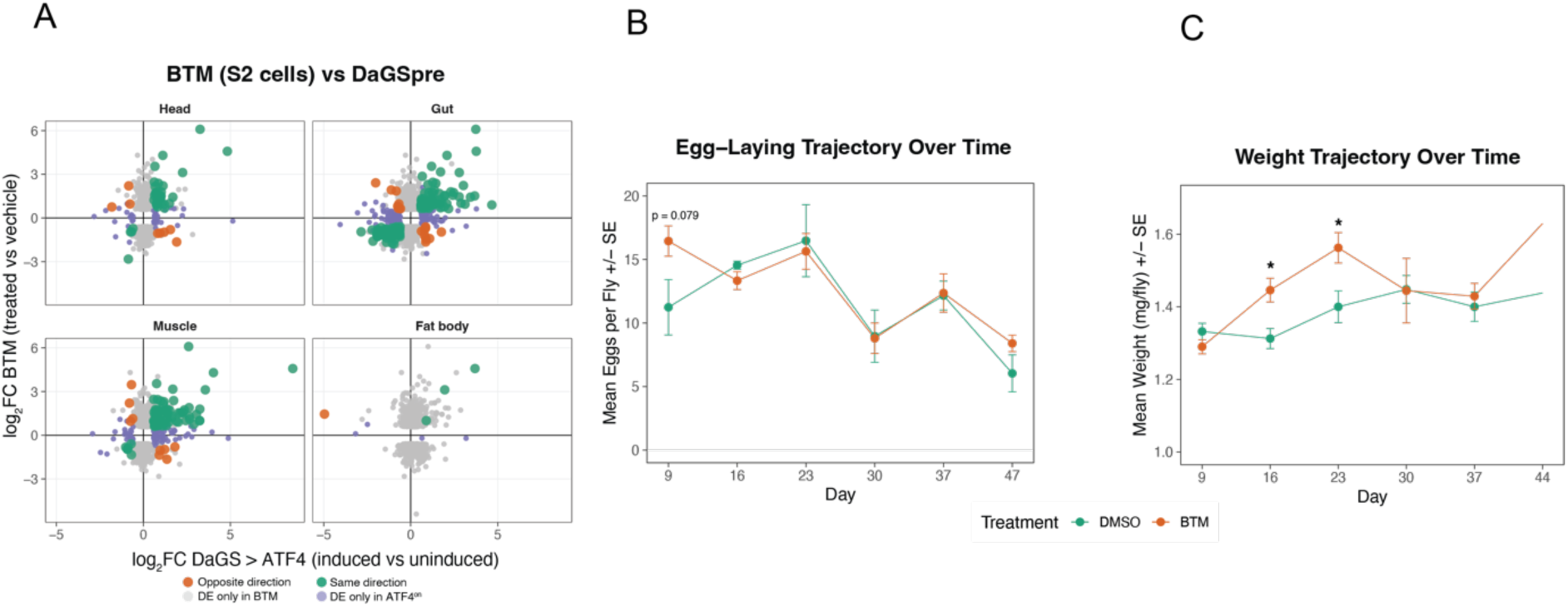
**a)** Comparison of the transcriptional response of Drosophila S2R+ cells to 10 µM BTM for 6 h with the response of ubiquitous ATF4^on^ flies during induction, in each tissue. Each point is a gene differentially expressed in at least one dataset, restricted to genes tested in both. The x-axis is log2 fold change in induced vs uninduced flies, and the y-axis is log2 fold change in BTM-vs vehicle-treated cells. Colours indicate genes changed in the same direction, in opposite directions, only in BTM-treated cells or only in ATF4^on^ flies. The responses overlapped more than expected by chance in head, gut and muscle, but not in fat body (one-sided Fisher’s exact test, Benjamini-Hochberg adjusted). Overlap, concordance and correlation statistics are given in Table S5. **b)** Lifetime fecundity of flies fed 10 µM BTM compared with vehicle (DMSO) controls, measured as eggs per female per 24 h. Data shown as mean ± SE. n = 5 vials of 5 females per condition per timepoint. No significant differences were detected at any timepoint (day 9, p = 0.079), Welch’s t-test per day. **c)** Lifetime weight measurements of whole flies fed 10 µM BTM compared with DMSO controls. Data shown as mean ± SE. * Indicates p < 0.05, Welch’s t-test per day. n = 5-8 vials of 3-5 flies per experimental condition.

**Figure S7.**
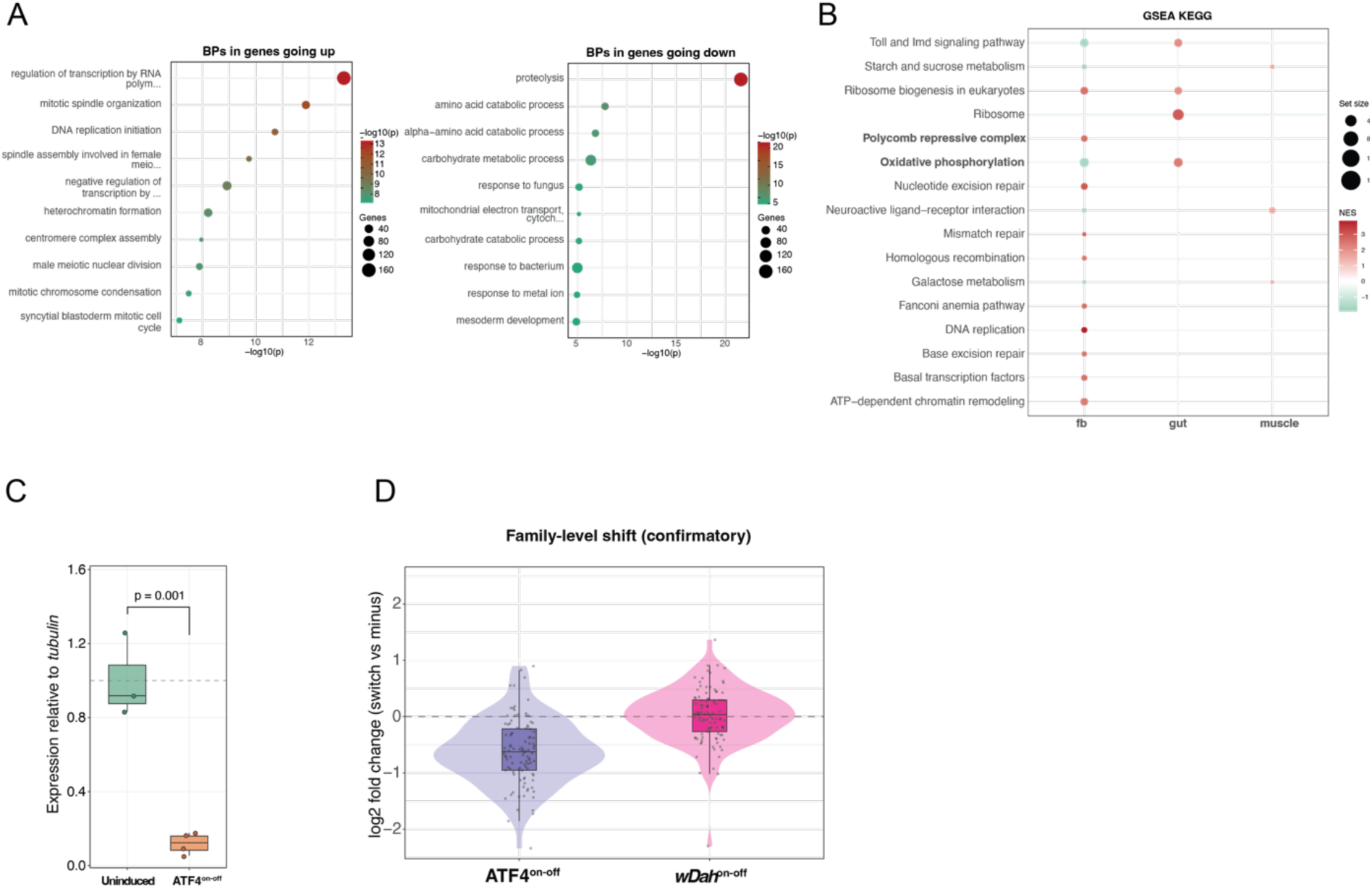
a) GO biological process terms enriched among genes up-(left) and downregulated (right) in the fat body of 50-day-old neuronal ATF4^on-off^ flies. Top 10 terms per direction among those at classic Fisher p < 0.05, ranked by the weight01 algorithm. b) Gene set enrichment analysis of KEGG pathways in fat body, gut and thoracic muscle of neuronal ATF4^on-off^ flies; no pathway reached significance in the head. Genes were ranked by ashrshrunk log2FC. Dot colour shows the normalised enrichment score (NES) and dot size the number of genes in each pathway. Only pathways significant at adjusted p < 0.05 are shown; Polycomb repressive complex and oxidative phosphorylation are highlighted. c) RT-qPCR quantification of *gypsy* transcripts in the fat body of 50-day-old neuronal ATF4^on-off^ and uninduced flies, normalised to *tubulin*. Box plots show the median and quartiles; points are individual pools of 12 fat bodies (n = 3–4 per condition). p = 0.001, Welch’s t-test on Log2 transformed values. **d)** Family-level shift in TE expression, confirmatory test. Violin plots show the distribution of per-family log2 fold changes (RU486 versus uninduced) in the fat body of 50-day-old neuronal ATF4^on-off^ flies and of *nSyb*-GS > *wDah* driver-alone controls; boxes indicate the median and interquartile range. The median family log2FC was compared against a null generated by exhaustively reassigning treatment labels between pools (ATF4^on-off^: median −0.63, p = 0.11, 35 arrangements, minimum attainable p = 0.029; *wDah*^on-off^: median +0.04, p = 0.93, 70 arrangements, minimum attainable p = 0.014).

**Figure S8.**
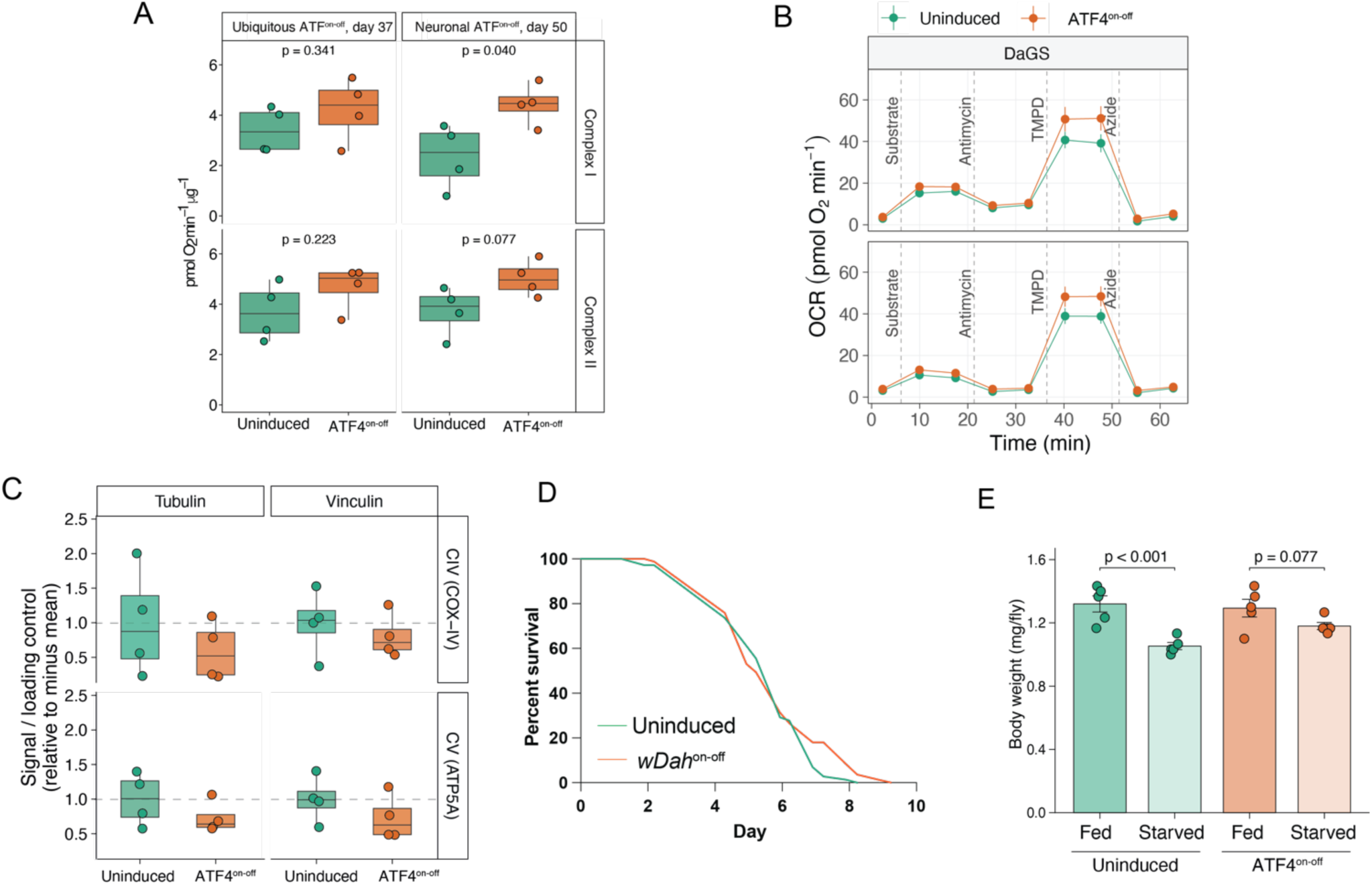
**a)** Respiratory chain capacity of fat body homogenates from ubiquitous, 35 day-old ATF4^on-off^ fat bodies, and 50 day-old neuronal ATF4^on-off^ flies compared with uninduced controls. Complex I capacity was measured with NADH and complex II capacity with succinate plus rotenone, both expressed as antimycin A-sensitive oxygen consumption normalised to protein (pmol O2 min^−1^ µg^−1^). Boxplots: quartiles; whiskers: extremes; overlay: individual cohorts. n = 4 cohorts per condition, technical wells averaged within each cohort. Ubiquitous: complex I p = 0.341, complex II p = 0.223. Neuronal: complex I p = 0.040, complex II p = 0.077, Welch’s t-test. **b)** Oxygen consumption traces of fat body homogenates from ubiquitous, 35 day-old ATF4^on-off^ fat bodies, compared with uninduced controls. Complex I (top) was fed NADH and complex II (bottom) was fed succinate plus rotenone, followed by sequential injections of antimycin A, TMPD/ascorbate and azide (dashed lines). Values are mean ± SEM of n = 4 cohorts, with technical wells averaged within each cohort. **c)** Western blot quantification of COX4 (complex IV) and ATP5A (complex V) in fat bodies of 50-day-old neuronal ATF4^on-off^ and uninduced flies, normalised to tubulin and vinculin. n = 4 per condition. p > 0.05, Welch’s t-test. **d)** Starvation survival (1% agarose) of 50-day-old *nSyb*-GS > *wDah* driver-alone flies. Uninduced n = 72 dead/5 censored, *wDah*^on-off^ n = 83 dead/1 censored; p = 0.24, log-rank test. **e)** Whole-body weight of 50-day-old neuronal ATF4^on-off^ flies compared to uninduced controls, fed or after 72h of starvation (1% agarose). Values are mean ± SEM of n = 5 samples of 3 flies per condition; dots show individual samples. Statistics are from a linear model on log-transformed body weight: brackets show fed versus starved within each condition (emmeans contrasts); RU:starvation interaction, ratio of fraction retained = 1.14, 95% CI 0.995-1.32, p = 0.058.

## Notes

https://www.ncbi.nlm.nih.gov/geo/query/acc.cgi?acc=GSE350078

